# Cellular and genomic properties of *Haloferax gibbonsii* LR2-5, the host of euryarchaeal virus HFTV1

**DOI:** 10.1101/2020.10.26.354720

**Authors:** Colin Tittes, Sabine Schwarzer, Friedhelm Pfeiffer, Mike Dyall-Smith, Marta Rodriguez-Franco, Hanna M. Oksanen, Tessa E.F. Quax

**Author notes:** These authors contributed equally (Colin Tittes, Sabine Schwarzer). **Correspondence should be addressed to**: Tessa E.F. Quax.

## Abstract

Hypersaline environments are the source of many viruses infecting different species of halophilic euryarchaea. Information on infection mechanisms of archaeal viruses is scarce, due to the lack of genetically accessible virus-host models. Recently a new archaeal siphovirus, Haloferax tailed virus 1 (HFTV1), was isolated together with its host belonging to the genus *Haloferax,* but it is not infectious on the widely used model euryarcheon *Hfx. volcanii.* To gain more insight into the biology of HFTV1 host strain LR2-5, we studied characteristics that might play a role in its virus susceptibility: growth-dependent motility, surface layer, filamentous surface structures and cell shape. Its genome sequence showed that LR2-5 is a new strain of *Hfx. gibbonsii.* LR2-5 lacks obvious viral defense systems, such as CRISPR-Cas, and the composition of its cell surface is different from *Hfx. volcanii,* which might explain the different viral host range. This work provides first deep insights into the relationship between the host of halovirus HFTV1 and other members of the genus *Haloferax*. Given the close relationship to the genetically accessible *Hfx. volcanii*, LR2-5 has high potential as a new model for virus-host studies in euryarchaea.

## INTRODUCTION

Viruses outnumber their microbial hosts by about a factor of 10 (Bergh *et al*., 1989; Wommack and Colwell, 2000; Suttle, 2007). Consequently, viruses have an important role in many ecosystems and impact microbial communities worldwide (Fuhrman, 1999; Suttle, 2007; Danovaro *et al*., 2016). Archaea are ubiquitous microorganisms that thrive both in extremophilic habitats, such as thermal hot springs and hypersaline lakes, as well as in mesophilic environments, like the oceans, soil and human gut (Karner *et al*., 2001; Lloyd *et al*., 2013; Lurie-Weinberger and Gophna, 2015). Many archaeal viruses differ significantly from those infecting bacteria and eukaryotes. The shapes of archaeal virions and the viral genomes are characterized by a high level of diversity. Some archaeal viruses, especially those infecting hyperthermophilic crenarchaea, have unique morphologies that are not encountered for viruses infecting bacteria and eukaryotes (Prangishvili *et al*., 2017; Munson-Mcgee *et al*., 2018). Other archaeal viruses, mainly infecting euryarchaea, display morphologies shared with some bacterial viruses (bacteriophages), such as head-tail or icosahedral shapes (Pietilä *et al*., 2014; Prangishvili *et al*., 2017). The study of archaeal viruses has been important to gain insight into the origin and evolution of viruses in general (Forterre and Prangishvili, 2009). The large majority of genes carried by archaeal virus genomes encode proteins of unknown function and consequently many aspects of the interaction between these viruses and their hosts remain enigmatic (Prangishvili *et al*., 2017; Krupovic *et al*., 2018). Studies on infection mechanisms or host recognition complexes of archaeal viruses are rare, but have yielded surprising results showing that infection strategies can be unique or display similarities with viruses infecting other domains of life (El Omari *et al*., 2019; Santos-Pérez *et al*., 2019). For example, entry and egress mxechanisms of archaeal viruses can rely on fusion with and budding of the virus through the cell membrane, respectively, as observed for eukaryotic viruses, or on the formation of archaeal specific pyramidal egress structures (Bize *et al*., 2009; Snyder *et al*., 2013; Quemin *et al*., 2016; El Omari *et al*., 2019). To advance studies on virus host interactions and infection mechanisms in archaea, model systems consisting of a well characterized host and virus are needed. Halophilic euryarchaea have proven a rich source of archaeal viruses. To date, more than 100 haloarchaeal viruses have been isolated of which the majority are tailed icosahedral double-stranded DNA viruses representing the order *Caudovirales* (Atanasova, Bamford, *et al*., 2015). Several haloarchaea are excellent research organisms as they are straightforward to cultivate, have relatively fast doubling times, and for several of them elaborate genetic tools are available (Leigh *et al*., 2011; Cheng *et al*., 2017). *Haloferax volcanii* is widely used as genetically accessible model organism, and different aspects of its biology such as replication, cell division, protein turn-over, transcription, translation and defense against viruses have been studied in detail (Eichler and Maupin-Furlow, 2013; Hawkins *et al*., 2013; Duggin *et al*., 2015; Maier *et al*., 2015; Pohlschroder and Schulze, 2019; Haque *et al*., 2020; Schulze *et al*., 2020). Curiously, in contrast to other haloarchaeal genera, viruses for the genus of *Haloferax* are extremely rare (Atanasova *et al*., 2012; Atanasova, Demina, *et al*., 2015). The first reports of *Haloferax* infecting viruses are those of HF1 virus infecting *Haloferax lucentense* and *Hfx. volcanii*, and a defective provirus of *Hfx. mediterranei,* both of which are no longer available (Nuttall and Smith, 1993; Li *et al*., 2013), [M.L. Dyall-Smith, pers. Communication]. Recently a new virus was isolated infecting a member of the genus *Haloferax*. *Haloferax* tailed virus 1 (HFTV1) and its host, *Haloferax* sp. LR2-5, originate from the saline Lake Retba near Dakar in Senegal (Mizuno *et al*., 2019).

The siphovirus HFTV1 has an icosahedral head of ~50 nm diameter and a long non-contractile tail of ~60 nm (Mizuno *et al*., 2019). Four major protein types were detected in the HFTV1 virion. The linear, circularly permuted dsDNA genome of 38 kb encodes 70 ORFs, of which half have homology to haloarchaeal viral genes, such as the archaeal siphovirus HRTV-4 isolated from Margherita di Savoia, Italy, and of uncultivated haloviruses from the solar saltern Santa Pola, Spain (Mizuno *et al*., 2019). The genome is likely subjected to a headful packaging mechanism initiated from a *pac* site (Mizuno *et al*., 2019). HFTV1 has a narrow host range among the Lake Retba archaeal strains tested. Besides its original host *Hfx. sp.* LR2-5, it infects *Halorubrum sp.* LR1-23 but not any of the endogenous *Haloferax* strains (Mizuno *et al*., 2019).

Currently, 19 species of the genus *Haloferax* are listed in the NCBI taxonomy. Twentyeight complete or draft genome sequences from the genus *Haloferax* are available (Lynch *et al*., 2012; Becker *et al*., 2014) including the complete sequences of *Hfx. gibbonsii* strain ARA6 (Pinto *et al*., 2015), *Hfx. volcanii* strain DS2^T^ (Hartman *et al*., 2010), and *Hfx. mediterranei* strain R-4 (ATCC 33500) (Han *et al*., 2012).

To gain insight into the question why *Haloferax* strains seems so resilient to viral infection and what makes the LR2-5 strain an exception, we sequenced and annotated its genome and compared it to closely related strains. We could assign this isolate to the species *Hfx. gibbonsii* using a genome based taxonomy (TYGS) and consequently LR2-5 was renamed as *Hfx. gibbonsii* LR2-5. In addition, we studied its biological properties, such as growth, cell shape, motility, and composition of its cell surface, which will contribute to developing the LR2-5 and HFTV1 system as an attractive model for archaeal virus-host studies.

## RESULTS AND DISCUSSION

### Growth properties and cell -shape transition of *Hfx. gibbonsii* LR2-5 during growth

To gain insight into the optimal cultivation conditions of the environmentally isolated strain *Hfx. gibbonsii* LR2-5, growth media with different composition and several incubation temperatures were tested (Figure S1). The shortest doubling time of 3.5 hours was achieved when the strain was aerobically grown in rich YPC medium with a total salt concentration of 18 % (wt/vol) at 42 °C (Figure S1). The doubling time of cells grown at 37 °C is about 4.5 hours in MGM medium and 6.5 hours in CA medium. Cultures grown at 37 °C exhibit prolonged lag-phases compared to cells grown at higher temperatures whereas cultures grown at 45 °C reach lower final optical densities (Figure S1).

Within the tested range (18 % and 23 %), the salt concentrations had no effect on the doubling times. The highest final optical densities under all conditions were reported in YPC and CAB medium with 18 % (wt/vol) salt concentration. The optimal growth medium and conditions are similar to those of *Hfx. volcanii* H26 (a derivative of strain DS2^T^, see supplementary text) (Jantzer *et al*., 2011), facilitating use of methods established for this model strain.

*Hfx volcanii* is reported to change its shape during growth in CA medium (Li *et al.*, 2019; Silva *et al*., 2020). We analyzed the cell shape of LR2-5 in CA medium by phase contrast light microscopy. This revealed that the cells were rod-shaped and usually ca. 1.5 – 4 μm in length in early-exponential growth phase (below OD_600_ 0.2) (Table S1). As the cultures reached mid-exponential growth phase (OD_600_ 0.2 to 0.6), a mix of rod- and plate-shaped cells was observed. In stationary phase (above OD_600_ 0.6), all cells appeared plate-shaped (Figure 1A). This transition from rod- to plate-shaped cells is reminiscent of what was recently observed for *Hfx. volcanii* (Li *et al*., 2019; Silva *et al*., 2020). However, whereas *Hfx. volcanii* is rod-shaped only during the very early exponential growth phase (below OD_600_ 0.1), *Hfx. gibbonsii* LR2-5 maintains the rod-shape much longer during development. The differences between the strains might originate from the longer adaptation of *Hfx. volcanii* to laboratory culture conditions where motility is not an evolutionary advantage, as the rod-shape seems to be linked with motility in *Hfx. volcanii* (Duggin *et al*., 2015; Li *et al*., 2019).

**Figure 1:**
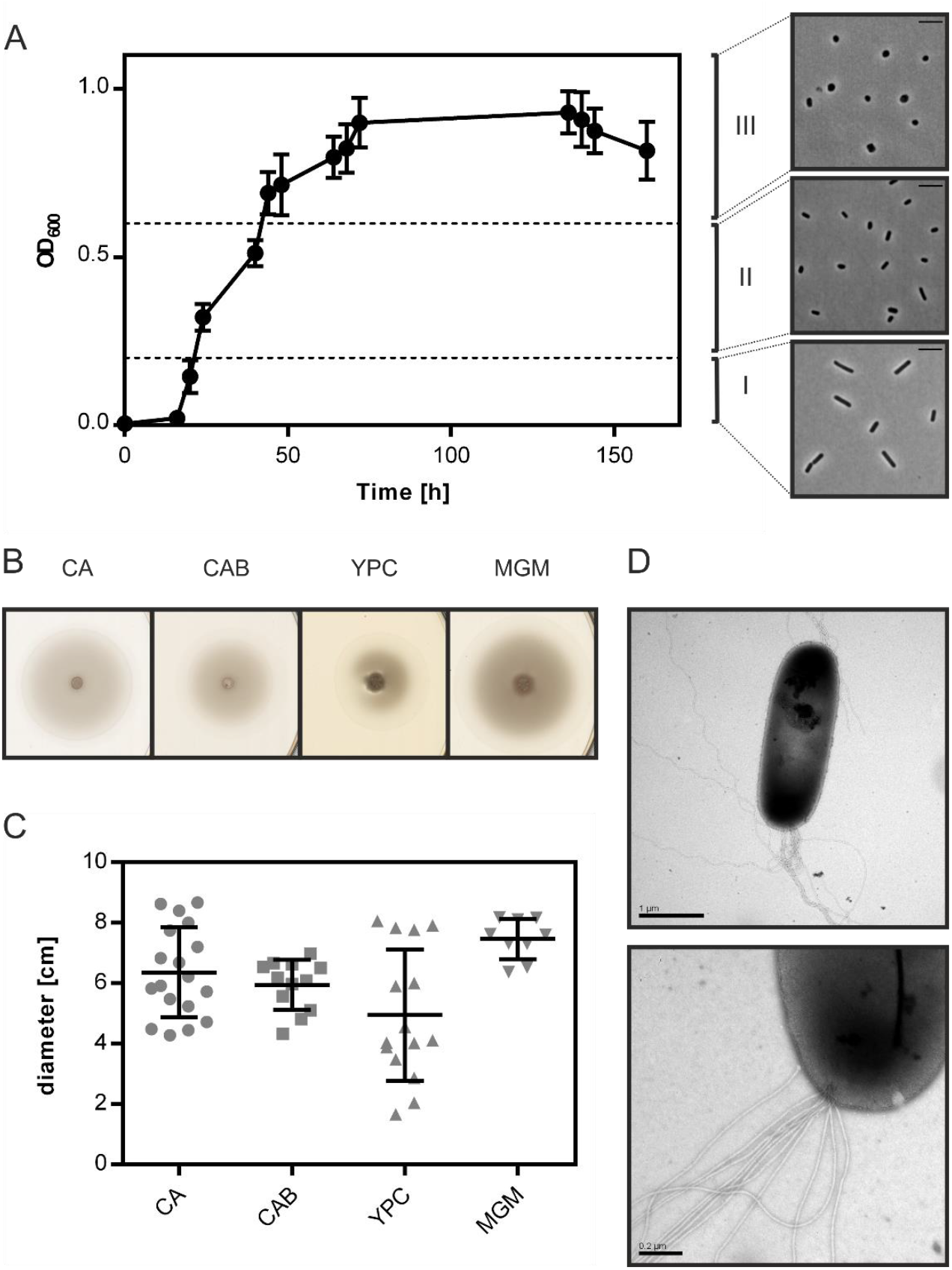
Cell shape change and motility of *Hfx. gibbonsii* LR2-5. **A:** Left side: typical growth curve of *Hfx. gibbonsii* LR2-5 in CA medium with 18 % (wt/vol) SW at 42 °C. Average optical density at 600 nm (OD_600_) was calculated from three independent technical replicates, error bars represent the standard deviation. Right side: phase contrast images show typical cell shapes correlating with three different growth phases: early-exponential (I) mid-exponential (II) and stationary (III) growth phases. **B:** Representative example of motility rings from *Hfx. gibbonsii* LR2-5 on semi-solid agar plates with different media after 3 days growth at 45 °C. **C:** Quantification of the diameter of motility rings formed on semi-solid agar plates with different media in 18 % SW. Calculations were made using more than three independent experiments including three biological replicates each. Middle line indicates the mean, lower and upper lines the standard deviation. **D:** Cells from early-exponential growth phase were negatively stained with 2 % (wt/vol) uranyl acetate. Top: *Hfx. gibbonsii* LR2-5 cell with typical rod-shaped morphology and archaella at the cell pole. Scale bar, 1 μm. Bottom: Close up of a bundle of archaella filaments at the cell surface. Scale bar, 0.2 μm.

### *Hfx. gibbonsii* LR2-5 is a motile archaeon

As *Hfx. gibbonsii* LR2-5 transitions from rod-to plate-shaped cells in a similar fashion as *Hfx. volcanii*, we tested the correlation of cell-shape with motility. First, the strain was stab inoculated on semi-solid agar plates. This showed that LR2-5 formed motility rings of ~6 cm diameter after 3 days in several different media (Figure 1B, C), indicative of an intact motility machinery and chemotaxis system.

Cells grown in liquid medium were observed with phase contrast microscopy at the optimal growth temperature of 42 °C. Time lapse imaging showed that cells displayed swimming motility in different growth media, of which an example is shown in (Movie S1). The average swimming speed was 6.5 μm/s. Analyzing the motility using time lapse microscopy during different growth phases showed that rod-shaped cells that are present during the early-exponential growth phase were highly motile, while the plate-shaped cells were immotile (Table S1). The motile phase of *Hfx. gibbonsii* LR2-5 is prolonged in comparison with that of *Hfx. volcanii*, correlating with the rod-shape appearance of the LR2-5 cells. The transition from motile rod-to immotile plate-shaped cells seems thus not confined only to *Hfx. volcanii*, but might be more general for members of the genus *Haloferax.*

Finally, *Hfx. gibbonsii* LR2-5 cells from early exponential phase were examined via transmission electron microscopy, revealing large bundles of 7 – 10 archaellar filaments per cell (Figure 1 D).

### Susceptibility of LR2-5 related *Haloferax* strains to HFTV1

The susceptibility of *Hfx. gibbonsii* LR2-5, *Hfx. gibbonsii* Ma2.38^T^ and *Hfx. volcanii* H26 (see supplementary text for information on strains) to HFTV1 was examined in parallel using a spot-on-lawn assay (Juez *et al*., 1986; Allers *et al*., 2004; Mizuno *et al*., 2019). Serial dilutions of the HFTV1 lysate (5 × 10^11^ PFU/mL) were spotted onto lawns of the three strains of *Haloferax* and the plates incubated for 3 days (Figure 2). Clearing of the cellular lawn appeared until a 10^−9^ dilution of virus lysate on the original virus host *Hfx. gibbonsii* LR2-5 (Figure 2 A), whereas only faint spots of hazy appearance were observed when undiluted HFTV1 lysate was spotted on *Hfx. gibbonsii* Ma2.38 (Figure 2 B). No zones of inhibition or separate plaques were observed when HFTV1 was spotted on *Hfx. volcanii* H26 (Figure 2 C). This was confirmed by traditional plaque assay. These results show that among the analyzed *Haloferax* strains, HFTV1 is only capable of efficiently infecting its original host *Hfx. gibbonsii* LR2-5. The absence of HFTV1 plaque formation on other strains is in line with previous results showing that HFTV1 is not infectious on six environmental *Haloferax* strains, isolated from Lake Retba (Mizuno *et al.*, 2019).

**Figure 2:**
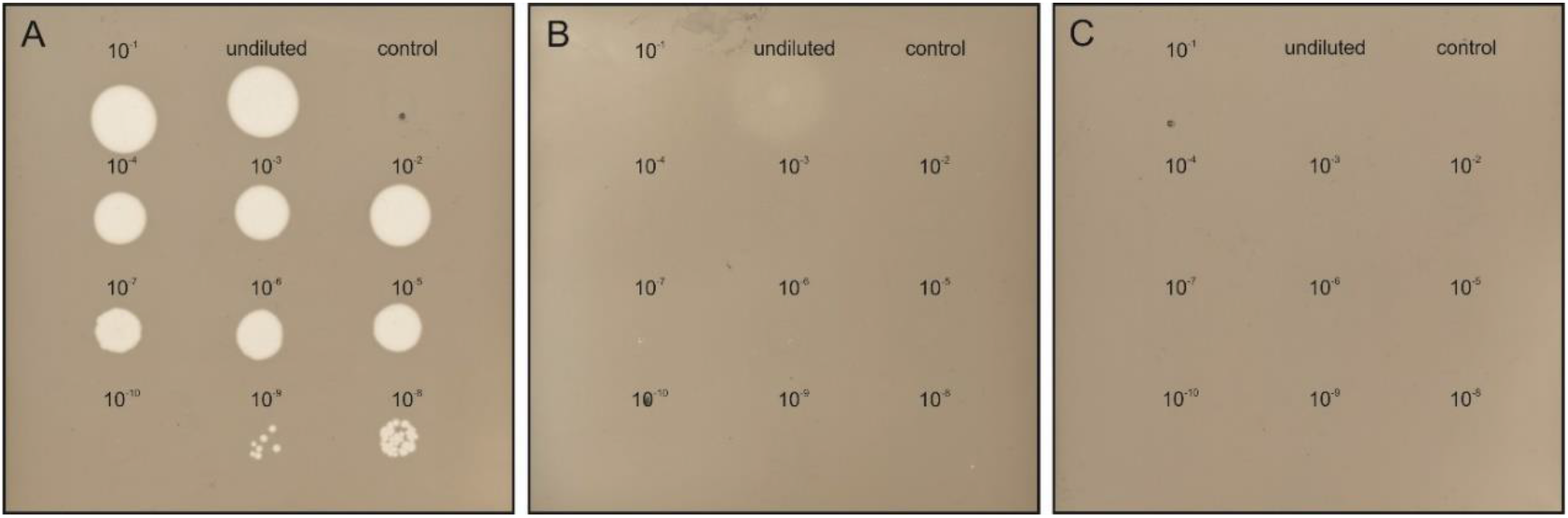
HFTV1 susceptibility of *Haloferax* strains by spot-on-lawn assay. Spot-on-lawn assay conducted with lawns of (**A**) *Hfx. gibbonsii* LR2-5, (**B**) *Hfx. gibbonsii* Ma2.38, and (**C**) *Haloferax volcanii* H26. Different dilutions of HFTV1 lysate (undiluted 5 × 10^11^ PFU/mL) were spotted on the host lawns and incubated for three days. Control spots were prepared with medium.

### *Hfx. gibbonsii* LR2-5 has a main chromosome and three plasmids

To sequence the genome of *Hfx. gibbonsii* LR2-5, DNA was extracted from exponentially growing *Hfx. gibbonsii* LR2-5 cells. PacBio genome sequencing and automated assembly of the sequences resulted in four circular contigs representing the four circular replicons (chromosome and three plasmids). Key characteristics of the genome are given in Table 1, and a more extensive summary is provided in Table S2.

**Table 1.** Replicons of *Hfx. gibbonsii* LR2-5. Each rRNA operon has 3 rRNAs (16S, 23S, 5S). Non-coding RNA (ncRNAs) are the 7S RNA, RNAse P RNA and the H/ACA guide RNA. Copy number of the replicon estimated by ratio of illumina read coverage (average) of each replicon relative to the average read coverage of the chromosome.

| *Replicon* | *Replicons* |  |  |  |
| --- | --- | --- | --- | --- |
|  | **Chromosome** | **pHGLR1** | **pHGLR2** | **pHGLR3** |
| *Length (bp)* | 2,999,641 | 608,598 | 322,970 | 65,035 |
| *GC (%)* | 66.9 | 61.7 | 66.9 | 55.8 |
| *Proteins (total)* | 3116 | 579 | 301 | 68 |
| *Number of rRNA operons* | 2 | - | - | - |
| *Number of ncRNAs* | 3 | - | - | - |
| *Number of tRNAs* | 53 | 1 | 1 | 0 |
| *Relative Copy Number* | (1.00) | 1.01 | 0.81 | 2.38 |

All replicons had at least 115-fold mean coverage. Illumina sequencing was used to validate and, if required, correct the assembled genome sequence, as described in the methods. This followed an established procedure (Pfeiffer *et al.*, 2020).

First, the strand and the point of ring opening was selected. For the chromosome, we adopted the convention of choosing a start position close to a canonical replication origin. However, we used a biologically relevant variation that we have used previously for *Natronomonas moolapensis* (Dyall-Smith *et al*., 2013), *Halobacterium hubeiense* (Jaakkola *et al*., 2016) and *Hbt. salinarum* strain 91-R6^T^ (Pfeiffer *et al*., 2019, 2020). This point of ring opening highlights the strong syntenic association of the genes adjoining the major replication origin. On one side is a highly conserved Orc/Cdc6 family member (gene *orc1* for strain LR2-5, see Table S3). On the other side, on the opposite strand, is the *oapABC* cluster (oap: origin-associated protein) (Wolters *et al*., 2019). For the plasmids, the applied strategy is detailed in the supplementary material.

The genome sequence was used to assign the LR2-5 strain to a taxon using the Type Strain Genome Server (TYGS; https://tygs.dsmz.de) (Meier-Kolthoff and Göker, 2019). It was classified within *Haloferax gibbonsii* with 90.6% average branch support. When using the complete genome including plasmids, the average branch support increased to 92.3%. The strain is now designated *Haloferax gibbonsii* LR2-5. Issues relating to the “16S rRNA based assignment” of this strain by the TYGS are described in the supplementary information.

### Comparative features and integrative elements of the replicons

The genome size and GC% are similar to the average values for other sequenced members of the genus *Haloferax* (3.8 Mb, 65.4% G+C; https://www.ncbi.nlm.nih.gov/genome/15284). Two of the three plasmids (pHGLR1 and pHGLR3) show significantly lower average GC (%) compared to the main chromosome. Tetranucleotide analysis (see methods) revealed that the motif CTAG and its inverse GATC, which are usually common in haloarchaeal genomes, are significantly under-represented in the LR2-5 genome, particularly on the main chromosome (0.19 odds ratio for both) and plasmid pHGLR2 (odds ratios 0.12 and 0.18). Methylated bases (^m4^C and ^m6^A) were detected within four distinct sequence motifs (Table S4). Only one of the four motifs (CTAG) was palindromic. The most frequently modified motif (GCG ^m4^CTG) was methylated on only one strand while the other three were methylated on both strands.

We compared the *Hfx. gibbonsii* LR2-5 replicons to those of the closely related model strain *Hfx. volcanii* DS2^T^ (see supplementary text) as well as *Hfx. gibbonsii* strains ARA6 and MA2.38^T^. For strain ARA6, a complete genome sequence is available. For type strain Ma2.38^T^, only a draft genome is available, consisting of contigs with unresolved replicon structure. However, this strain is available in culture collections and could be used for virus susceptibility tests described above. BLASTn comparisons show a close similarity of the *Hfx. gibbonsii* LR2-5 chromosome to those of *Hfx. gibbonsii* strains Ma2.38^T^ and ARA6 (the red and green BLASTn rings, Figure 3A), and to *Hfx. volcanii* DS2^T^ (orange ring). Regions of sequence variation between strains (blank sections of the BLASTn rings) are frequently also regions of lower than average GC (troughs in the GC content plot) and predicted genomic islands (grey bars). These have been labeled either as IGEs (integrative genetic elements) or Long Variable Regions (LVR), and their details are given in Table S5.

**Figure 3.**
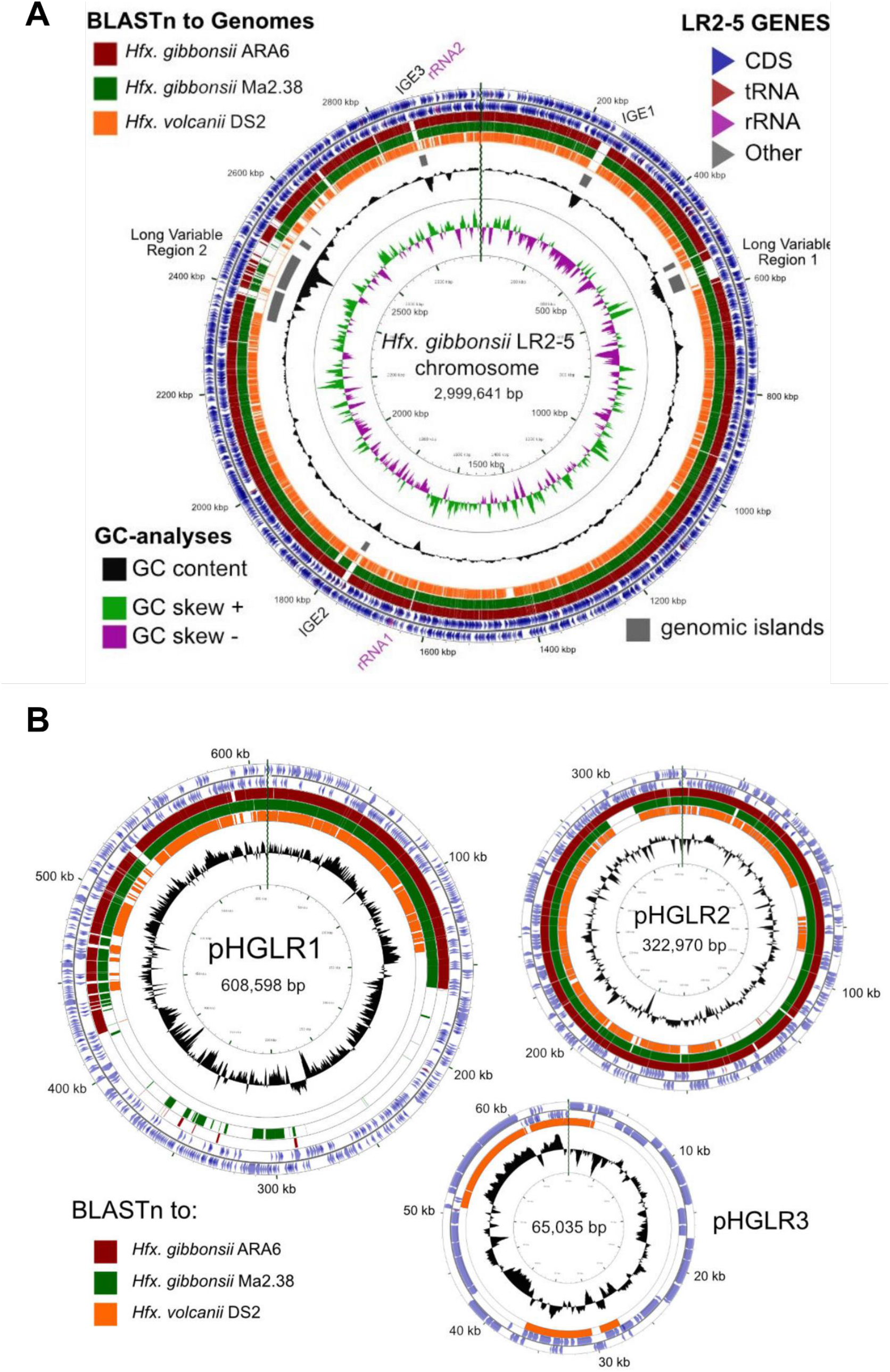
Comparison of the *Hfx. gibbonsii* LR2-5 chromosome and plasmids to those of three close relatives. **A:** *Hfx. gibbonsii* LR2-5 chromosome map showing similarity to replicons of closely related strains. The two outermost rings depict the annotated genes (CDS, tRNA and rRNA) of strain LR2-5, for the forward and reverse DNA strands (color key, upper right). Rings 3-5 depict BLASTn similarities between strain LR2-5 and the three strains listed in the color key (upper left). Colored bars represent regions of similarity (Expect value ≤ 10^-20^, cutoff=90% nucleotide identity) while uncolored (white) regions represent no significant similarity. Ring 6 (grey blocks) are predicted genomic islands (IslandViewer 4). The two innermost rings represent plots of GC content (black) and GC-skew (green/purple) for strain LR2-5, and the color key for these plots is given in the lower left. The GC content ring plots differences from the average GC%, with outwards pointing peaks indicating higher than average, and inwards pointing peaks indicating lower than average GC%. IGE1-3, integrative genetic elements (see text). rRNA1 and rRNA2 are the two ribosomal RNA operons. Tick marks around the inner-most and outermost rings show DNA size in kb. The maps and plots were made using the CGView Server (http://stothard.afns.ualberta.ca/cgview_server). **B:** *Hfx. gibbonsii* LR2-5 plasmid maps showing similarity to replicons of closely related strains. The color key for LR2-5 genes (outer two rings) and the inner GC content plots (black) are as described in **A**. The comparison strains used to produce the BLASTn similarity rings are indicated by the colored boxes lower left. The actual replicons used are as follows. Plasmid pHGLR1 is compared to *Hfx. gibbonsii* ARA6 plasmid pHG1 (488,062 bp) (red), *Hfx. gibbonsii* Ma2.38 (green) and to *Hfx. volcanii* DS2^T^ plasmid pHV3 (437,906 bp) (orange). Plasmid pHGLR2 is compared to *Hfx. gibbonsii* ARA6 plasmid pHG2 (335,881 bp) (red), contig 28 (AOLJ01000028) of *Hfx. gibbonsii* Ma2.38 (green), and to *Hfx. volcanii* DS2^T^ plasmid pHV4 (635,786 bp) (orange). Plasmid pHGLR3 is compared to *Hfx. volcanii* DS2^T^ plasmid pHV1 (85,092 bp) (orange).

The relation between the chromosomes of *Hfx.gibbonsii* LR2-5, *Hfx. gibbonsii* ARA6 and *Hfx. volcanii* DS2^T^ has been additionally analyzed by MUMmer, presented as dotplots (Figure S2). Both plots show a long inversion of the LR2-5 genome relative to the others, with the inversion boundaries of LR2-5 being the two inward facing rRNA operons (nt 1658979-1663984 and nt 2916773-2921778). This inversion correlates with the abrupt shifts in the GC skew which occur close to the rRNA operons (Figure 3A).

All three IGEs have integrated into tRNA genes on the chromosome, have terminal direct repeats partially duplicating the 3’ part of the tRNA, and carry a gene for XerC/D integrase near the end which is adjacent to the complete copy of the tRNA. IGEs could be provirus-related. IGE3 shows strong similarity to a betapleolipovirus-like provirus of *Hfx. prahovense* Arc-Hr and to betapleolipoviruses HRPV9 (Atanasova *et al.*, 2018) and HGPV-1(Senčilo *et al*., 2012) of the family *Pleolipoviridae* (Demina and Oksanen, 2020)(Figure S3), although IGE3 appears to have lost many viral core genes. The other two IGEs are less clear but may also represent provirus-remnants. IGE1 carries three genes for restriction-modification proteins, as well as a gene (HfgLR_01385) that is commonly found in proviruses reminiscent of the *Caudovirales* type viruses of other haloarchaea, e.g. *Haloferax* sp. ATB1 (accession JPES01000032).

The long variable regions LVR1 and LVR2 represent longer and more complex regions of variability than IGEs. They seem hotspots of recombination and their borders could not be easily defined. LVR1 carries replication genes *(orc6, polB2)* and may be plasmid related. A comparison between *Hfx. gibbonsii* LR2-5 and ARA6 across the 38 kb of LVR1 revealed the marked differences in gene composition and size (Figure S4). LVR2 is almost 195 kb in length and also carries replication genes (e.g. *orc10, orc11)* as well as many genes that could influence virus susceptibility of the host, including the genes encoding PilA2, AglF, AglM, and many enzymes involved in sugar metabolism (glycosyltransferases, D-galactonate dehydratase, sugar-nucleotidyltransferases) as well as secreted glycoproteins.

All three plasmids match to plasmids of *Hfx. volcanii* DS2^T^ (Figure 3 B, orange rings). Plasmids pHGLR1 and pHGLR2 show strong nucleotide similarity to plasmids of *Hfx. gibbonsii* strains ARA6 and to contigs of strain Ma2.38^T^ (Figure 3B; red and green BLASTn rings). Plasmid pHGLR3 does not have a counterpart in the other *Hfx. gibbonsii* strains. All three plasmids also have strain specific parts. The strain specific parts of pHGLR1 and pHGLR3 tend to have lower than average GC content, and this is particularly evident in the 150-420 kb region of pHGLR1 (56.5% G+C) compared to the rest of this plasmid (66% G+C), a difference of almost 10 percentage points (Figure 3B). This may indicate that pHGLR1 is the result of a large integration or fusion event. A similar example has been described previously in the *Hbt. salinarum* plasmid pHS3, which carries a 70 kb high-GC island (Barylski *et al*., 2020). The borders of these two regions share a similar gene cluster of four ABC-transport genes (*tsg*) that show significant nucleotide similarity, e.g. *tsgD5* (HfgLR_20645; HVO_A0145) and *tsgD6* (HfgLR_22070; HVO_A0281), but are outwardly oriented relative to each other.

Plasmid pHGLR2 has an 11.6 kb stretch of DNA near the 300 kb mark (nt 291722 – 303338; HfgLR_24335 to HfgLR_24410) that is shared only with pHG2 of strain ARA6, and is absent in strain Ma2.38^T^ and *Hfx. volcanii* DS2^T^ (pHV4). This region of pHGLR2 carries genes for various transporters, phosphoribosyl-AMP cyclohydrolase, and several uncharacterized proteins.

*Hfx. gibbonsii* LR2-5 has two rRNA operons and 55 regular tRNA genes, of which 53 are encoded on the main chromosome (Table 1). In addition, there are 11 partial tRNAs, of which 9 are on the chromosome. In several cases, the remnant is directly adjacent to a full copy of the same tRNA. Each of the three IGEs has duplicated termini, one being the tRNA and the other a tRNA remnant. For more information see supplementary material.

A detailed analysis of the transposons (Table S6) revealed that *Hfx. gibbonsii* strains ARA6 and LR2-5 have a low number of transposons and other mobile genetic elements compared to other haloarchaea (12 in strain ARA6, 31 in strain LR2-5; Table S6). For more information see supplementary text.

### Comparison of protein coding genes of LR2-5 and HFTV1-resistant *Haloferax* species

Overall, 3204 of the 4064 proteins (~79%) encoded in the *Hfx. gibbonsii* LR2-5 genome have an ortholog in *Hfx. volcanii* DS2^T^ with an average of 93 % protein sequence identity. One-third (37 %) of these orthologs have 96 – 98 % protein sequence identity. The distribution is slightly uneven between the chromosome (84% of proteins have an ortholog) and the plasmids (61% of proteins have an ortholog).

To gain insight into the differences between *Hfx. gibbonsii* LR2-5 and the other *Haloferax* strains, we compared protein coding genes between these species with a focus on anti-viral defense systems and genes encoding possible viral anchor points and receptors at the cell surface (Tables S6-S11). We considered *Haloferax* strains for which a complete genome sequence is available, *Hfx. volcanii* DS2 and *Hfx. gibbonsii* ARA6.

### Anti-viral defense systems

The arms race of bacteria and archaea with their viruses has led to the development of a plethora of defense mechanisms against viruses. These include CRISPR-Cas, toxin antitoxin (TA) systems, Restriction Modification (RM) systems, and several recently discovered new systems (Stern and Sorek, 2011; Azam and Tanji, 2019).

In bacteria, TA systems are sometimes part of an antiviral defense mechanism relying on abortive infection (Gerdes *et al*., 2005; Tachdjian and Kelly, 2006). *Hfx. gibbonsii* LR2-5 encodes a few TA systems (Table S7), although none of the functional systems appear likely to be linked with abortive infection. Likewise, RM systems can play a role in defense against foreign genetic elements in bacteria (Tock and Dryden, 2005). Several type I RM systems were predicted in the LR2-5 genome which are only marginally related to *Hfx. volcanii* RM systems (Table S8, supplementary text). This, substantiated by the differences in DNA methylation (Table S3, rebase data: http://rebase.neb.com/cgi-bin/pacbioget?5891), indicates that RM systems may be an infection barrier in *Hfx. volcanii* and *Hfx. gibbonsii* MA2.38. Indeed, the HFTV1 genome contains 24 target sites for the *Hfx. volcanii* Mrr restriction endonuclease.

Surprisingly, *Hfx. gibbonsii* LR2-5 does not contain any CRISPR-Cas systems. These anti-viral defense systems are very common among archaea and *Hfx. volcanii* contains a well-studied and functional CRISPR-Cas system (Maier *et al*., 2019). *Haloferax gibbonsii* MA2.38 is predicted to contain two CRISPR-Cas systems, as well as an additional CRISPR array (analyzed using https://crisprcas.i2bc.paris-saclay.fr/CrisprCasFinder/Index). Neither *Hfx. volcanii* nor *Hfx. gibbonsii* MA2.38 have any spacers matching the HFTV1 genome. On the other hand, *Hfx. gibbonsii* ARA6 does not contain any CRISPR-Cas systems. The lack of CRIPSR-Cas systems in LR2-5 does however make it an attractive model organism for the study of virus-host interactions as it is more likely to be susceptible to other viruses. Further details on defense systems can be found in the supplementary text.

### S-layer

The surface layer (S-layer) functions as a cell wall in many archaea and some bacteria and is also an attractive target for viruses at the cell surface. S-layer proteins are highly abundant and can display marked differences between strains and species. Several bacterial phages require host S-layer for infection (Mescher and Strominger, 1977; Edwards and Smit, 1991; Plaut *et al*., 2014). Archaeal S-layer proteins are usually heavily glycosylated (Albers and Meyer, 2011; Kaminski, Lurie-Weinberger, *et al*., 2013; Kandiba and Eichler, 2014).

To determine the S-layer glycoprotein (SLG) of LR2-5, we analyzed the total cell lysates of different *Haloferax* strains by SDS-gel electrophoresis (Figure 4A). In several archaea, the SLG is the most abundant cellular protein. In halophilic archaea it is easily detected by Coomassie staining as a prominent protein band running at ~ 245 kDa. A prominent band in LR2-5 cell lysate migrated slightly above that of the *Hfx. volcanii* SLG but identically to an equally prominent band of the type strain of *Hfx. gibbonsii* (MA2.38). The band from LR2-5 was excised and subjected to mass spectrometry analysis (data not shown). This showed the presence of HfgLR_04635 and HfgLR_11210 in this protein band. Both proteins are predicted to have similar molecular weights of ~90-95 kDa, but due to their abundant predicted glycosylation sites, they run at a very different height, as is common for halophilic proteins (Shalev *et al*., 2018). The function of HfgLR_11210 is not clear. The detection of HfgLR_04635 is consistent with the recent identification of a homolog of HfgLR_04635 as the SLG of a *Hfx. gibbonsii* strain (ABY42_04395) (Shalev *et al*., 2018). The two proteins show 99% protein sequence identity, but are only very distantly related to the SLG of *Hfx. volcanii* (HVO 2072; 24% protein sequence identity) (Sumper *et al*., 1990).

**Figure 4.**
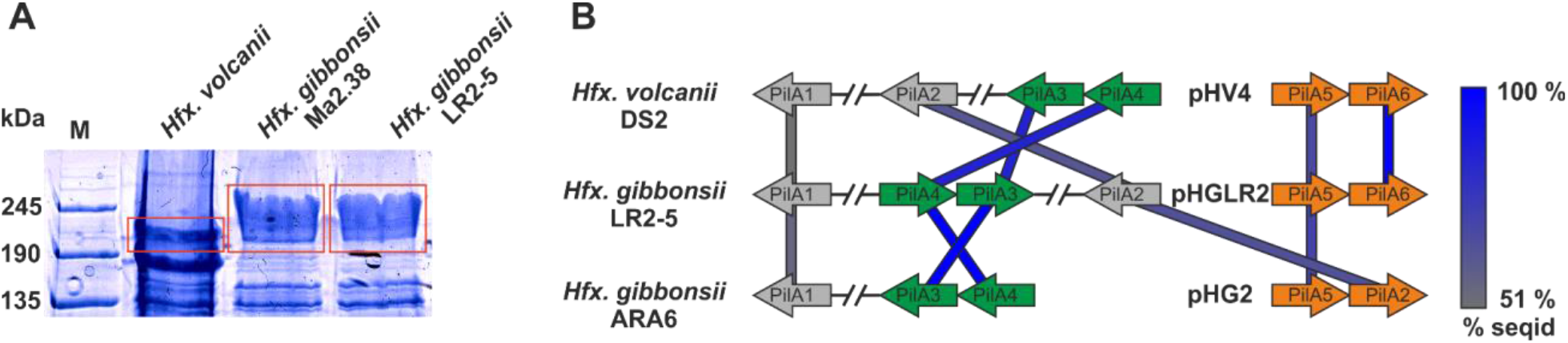
Surface protein and pili of *Hfx. gibbonsii* LR2-5. **A**: Coomassie stained SDS-PAGE gel of *Hfx. volcanii, Hfx. gibbonsii* Ma2.38 and *Hfx. gibbonsii* LR2-5 whole cell lysates. The prominent bands at around 245 kDa are considered to run at the height of the S-layer glycoproteins (red boxes). The LR2-5 band was subjected to mass spectrometry. **B:** Comparison of the genetic loci of *Hfx. gibbonsii* LR2-5 PilAs with those of *Hfx.gibbonsii* ARA6 and *Hfx. volcanii* DS2^T^. Homologs to all six *Hfx. volcanii* pilins are present in *Hfx. gibbonsii* LR2-5, however particularly PilA1, PilA2 and PilA5 are more distant homologs (protein sequence identity 51 – 68 %). PilA3 and PilA4 are organized in an operon in all three strains and the proteins have high protein sequence identity (≥ 85%).

### Pilins

Filamentous surface structures are well known primary anchor points for several viruses infecting bacteria (Poranen *et al*., 2002; Mäntynen *et al*., 2019). Recent data suggests that a few crenarchaeal viruses also tether to filamentous surface structures (Quemin *et al*., 2013; Hartman *et al*., 2019; Rowland *et al*., 2020). A hallmark of archaeal surface structures is the widespread similarity to bacterial type IV pili. Adhesive type IV pili of archaea are involved in attachment to biotic and abiotic surfaces which may lead to biofilm formation (Pohlschroder and Esquivel, 2015; van Wolferen *et al*., 2018). Pilins typically have a type III signal sequence, which is processed in *Haloferax* by the prepilin/prearchaellin peptidase PibD. After N-terminal processing, pilin subunits are inserted into surface filaments (Pohlschroder *et al*., 2018). The core membrane and biosynthesis complex for pilin subunit insertion is formed by the membrane platform protein PilC and the cytosolic assembly ATPase PilB (Pohlschroder *et al*., 2018). In *Hfx. volcanii*, six *pilBC* pairs are found.

While *pilB1C1* and its associated genes are present in *Hfx. gibbonsii* strain ARA6 but not in LR2-5, all other *pilBC* pairs with their associated genes are highly conserved between *Hfx. volcanii* strain DS2^T^ and *Hfx. gibbonsii* strains ARA6 and LR2-5 (Table S9).

*Hfx. gibbonsii* LR2-5 codes for 6 pilins, five of which are more closely related to those of *Hfx. gibbonsii* ARA6 than to those of *Hfx. volcanii* DS2^T^ (with PilA6 being absent from *Hfx. gibbonsii* ARA6, Figure 4B).

The individual roles in attachment and biofilm formation of the pilins of *Hfx. volcanii,* PilA1-6 seem to have slightly different functions in adhesion and microcolony formation (Esquivel *et al*., 2013; Legerme *et al*., 2016; Legerme and Pohlschroder, 2019).

The pilin genes are organized differently in the three *Haloferax* strains and some *Hfx. volcanii* pilins have only distant homologs in *Hfx. gibbonsii* LR2-5 and ARA6 (Figure 4A). Strikingly, *Hfx. gibbonsii* LR2-5 PilA1, PilA2 and PilA5 have closer homologs in other species, particularly *Hfx. sp.* Atlit-19N (isolated from high salt tide-pools on the coast of Israel (Atlit, summer 2012); BioProject PRJNA431124), than in *Hfx. volcanii* and *Hfx. gibbonsii* ARA6. As mentioned above, PilA2 is located within a long variable region in *Hfx. gibbonsii* LR2-5.

N-glycosylation plays a crucial role in pilus-mediated surface attachment and all *Hfx. volcanii* PilA pilins except PilA5 are glycosylated (Esquivel *et al*., 2016). All *Hfx. gibbonsii* LR2-5 PilAs except PilA2 are predicted to contain at least one N-glycosylation site (NxS/T) (Table S10). The predicted N-glycosylation sites in *Hfx. volcanii* and *Hfx. gibbonsii* LR2-5 are fully conserved for PilA3 but at maximum partially conserved for the other PilAs. Further details on type IV pili can be found in the supplementary text.

### Archaellum and chemotaxis machinery

The archaeal motility structure, the archaellum (archaeal flagellum), also displays homology to type IV pili (Albers and Jarrell, 2018). The archaellum is a rotating filamentous structure which functions analogously to the bacterial flagellum, as it provides swimming motility in liquid (Alam and Oesterhelt, 1984; Kinosita *et al*., 2016). However, archaella and flagella have a fundamentally different structural organization and their protein components are completely unrelated (Albers and Jarrell, 2015). Proteins related to archaella biogenesis and function are typically clustered in haloarchaeal genomes (Kalmokoff and Jarrell, 1991; Patenge *et al*., 2001; Jarrell and Albers, 2012). This *arl* cluster (previously *fla* cluster) is highly conserved between *Hfx. volcanii* and *Hfx. gibbonsii* strain LR2-5 (84 – 98 % protein sequence identity) with strictly conserved gene synteny.

The archaellum is involved in directional movement together with a bacterial-type chemotaxis system. The linking components have recently been identified (*cheF*, *arlCDE)* (Schlesner *et al*., 2009, 2012; Quax *et al*., 2018; Li *et al*., 2020). *Hfx. volcanii* and *Hfx. gibbonsii* LR2-5 have very similar *che* genes (89 – 100 % protein sequence identity). Both genomes show strictly conserved gene synteny in which the *che* gene cluster is split and surrounds the *arl* cluster. Conclusively, the genomic content of LR2-5 and the observed expression of archaella and motility machinery in liquid and semisolid agar (Figure 1B) show that LR2-5 is a highly motile strain. Due to the high level of conservation, the motility might not be responsible for the differences in virus susceptibility between strain LR2-5 and *Hfx. volcanii*.

### Protein N-glycosylation

N-glycosylation plays an important role in the biosynthesis and function of many surface exposed proteins such as pilins, archaellins and S-layer glycoproteins (Esquivel *et al*., 2016; Tamir and Eichler, 2017). N-glycosylation pathways differ between haloarchaeal species, and glycosylation of surface proteins, particularly the S-layer, may even vary depending on differing environmental factors such as salinity (Kaminski, Naparstek, *et al*., 2013). Indeed, *Hfx. volcanii* uses two distinct glycosylation pathways which were initially identified as being dependent on environmental salinity: the canonical pathway (AglB-J) and the low-salt pathway (Agl5-15) (Kaminski, Guan, *et al*., 2013). Unexpectedly, most genes coding for the low-salt N-glycosylation pathway were recently found to be expressed under optimal growth conditions (Schulze *et al.*, 2020).

The *Hfx. gibbonsii* LR2-5 glycosylation proteins are mostly found in two main chromosomal clusters (Figure 5A, B; marked I, II), with a few outliers, some being encoded on the plasmids (Figure 5C). Only 5 of the 20 characterized *Hfx. volcanii* Agl proteins have an ortholog in the LR2-5 genome (Table S11, Figure 5). These proteins appear to be well conserved within *Haloferax,* as close homologs are found in many other members of the genus. In addition, a number of LR2-5 genes in clusters I and II are distantly related to *Hfx. volcanii* glycosylation genes and thus are likely involved in N-glycosylation (Table S12, Figure 5).

**Figure 5.**
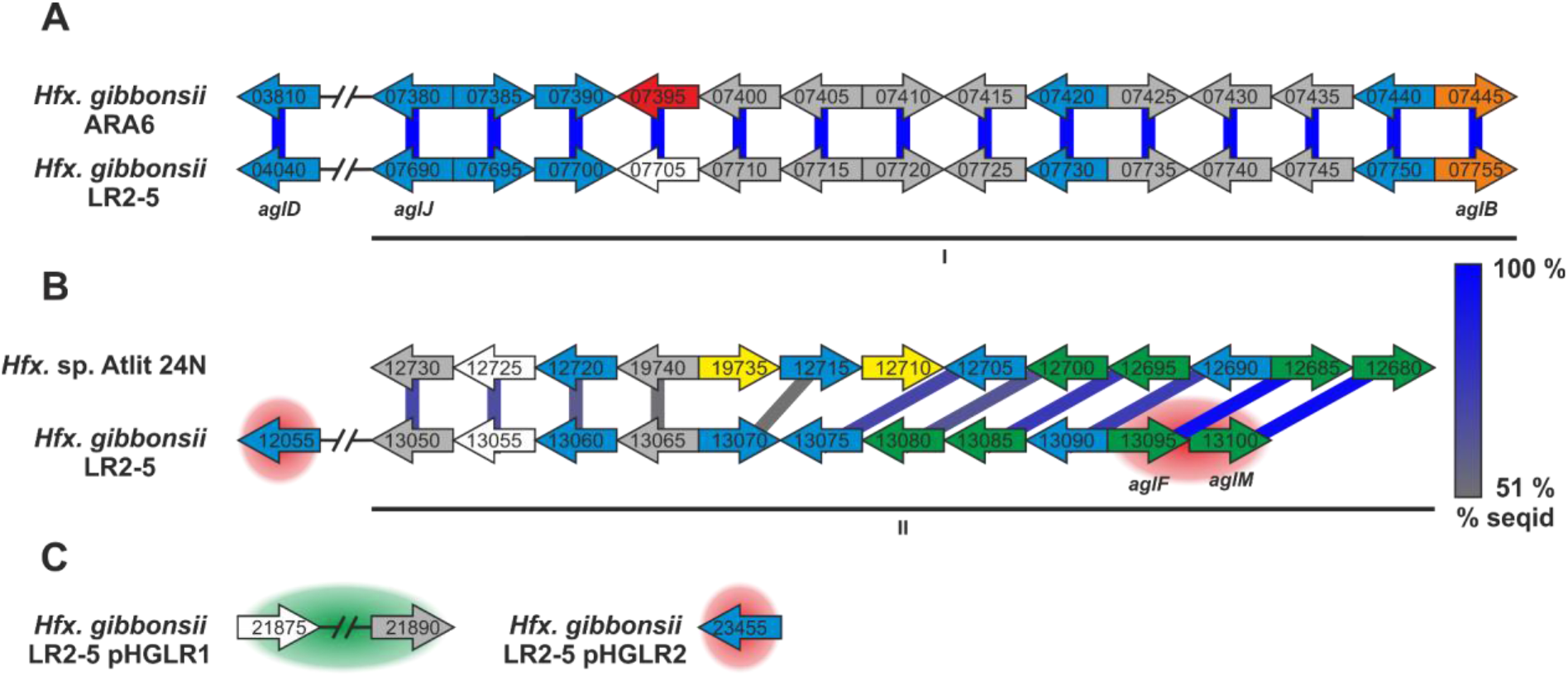
*Hfx. gibbonsii* LR2-5 N-glycosylation genes with comparison to their closest homologs. Genes are located in two clusters (marked as I and II in **A** and **B**) with some genes being located apart, e.g. on a plasmid (**C**). Close homologs in *Hfx. volcanii* are indicated with the corresponding *agl* name. Predicted protein function is indicated by color code; blue: glycosyltransferase, red: flippase, white: rfbX family protein (potential flippase), orange: oligosaccharyltransferase, green: sugar modification enzyme, grey: not directly related to N-glycosylation, yellow: hypothetical protein/pseudogene not apparently related to glycosylation. Protein homology and sequence identity is indicated by colored bars. Bar color indicates the level of amino acid sequence identity (see the color coding on the right). Background coloring indicates additional close homologs, red: *Hfx. gibbonsii* ARA6, green: *Hfx. sp.* Atlit-16N and Atlit-10N. Gene sizes are arbitrary.

Strikingly, the five genes with strong similarity to characterized *agl* genes of *Hfx. volcanii* are not in genomic vicinity, but spread across the two clusters with one outlier. The first cluster (Figure 5A, I) encodes proteins with close homologs in *Hfx. gibbonsii* ARA6, *Hfx. sp.* Atlit 4N, 6N, 10N, 16N and 19N as well as a few other *Haloferax* strains and species. The second cluster (Figure 5B, II) encodes proteins with closest homologs in *Hfx. sp.* Atlit 24N, 109R and 105R. Most of these proteins (particularly HfgLR_13050 through HfgLR_13070) have very few other close homologs. Like *pilA2,* this entire cluster lies within the long variable region LVR2.

Of the plasmid-encoded N-glycosylation proteins, HfgLR_23455 has many close homologs among proteins of other *Haloferax* species encoded also on plasmids, whereas HfgLR_21875 and 21890 have only four close homologs (> 70 % protein sequence identity, Table S12).

## CONCLUSION

We functionally characterized the euryarchaeon *Hfx. gibbonsii* LR2-5, the host of the siphovirus HFTV1, and sequenced its genome. *Hfx. gibbonsii* LR2-5 was shown to grow optimally under similar conditions as strains of the model species *Hfx. volcanii*, aiding in development of a genetic system for LR2-5. In addition, cells of *Hfx. gibbonsii* LR2-5 transition from motile rod-shaped cells to immotile plate-shape cells during growth. In comparison with the laboratory strain *Hfx. volcanii*, they stay rod-shaped much longer, facilitating studies of the motility machinery. Sequencing of the genome of *Hfx. gibbonsii* LR2-5 indicated that the cell surface of this strain likely differs from some of its close relatives. LR2-5 encodes several pilins that differ from those of *Hfx. volcanii* and *Hfx. gibbonsii* ARA6. Future work is required to analyze if the variability of these pilins is involved in the differences in virus susceptibility, as the receptor of HFTV1 has not been identified yet. The archaella of LR2-5 show high sequence conservation to other *Haloferax* species. Genes involved in glycosylation of surface proteins are very different between *Hfx. gibbonsii* and its close relatives. It is possible that the glycans of LR2-5 differ considerably from those of *Hfx. volcanii* and this difference might be responsible for the differences in virus susceptibility. Further analysis is needed to identify the actual glycans of LR2-5 and to establish if glycosylation is related to infection of HFTV1. Interestingly, *Hfx. gibbonsii* LR2-5 does not encode any CRISPR-Cas system, which is in contrast with the yet analyzed *Haloferax* strains that were found to be resistant to HFTV1. In addition, its R/M systems differ significantly from these other strains. Therefore, the absence of a CRISPR-Cas and the Mrr restriction endonuclease system might also explain the susceptibility of LR2-5 to HFTV1. This work will enable future research on HFTV1 adsorption to the host cell and interaction with anti-viral defense systems. Moreover, it significantly contributes to the development of *Hfx. gibbonsii* LR2-5 into a model system for the study of archaeal virus-host interactions.

## MATERIALS AND METHODS

### Archaeal strains and viruses, media and growth conditions

*Haloferax gibbonsii* LR2-5 (previous *Haloferax* sp. LR2-5; (Mizuno *et al*., 2019)), *Haloferax gibbonsii* MA2.38^T^ (Juez *et al*., 1986) and *Hfx. volcanii* H26 (Bitan-Banin *et al*., 2003) cells were cultured as described previously (Nuttall and Smith, 1993; Allers *et al*., 2004; Duggin *et al*., 2015; Mizuno *et al*., 2019). HFTV1 virus was grown and virus stocks were prepared as described (Mizuno *et al*., 2019). For details, see supplementary information.

### Transmission electron microscopy

*Hfx. gibbonsii* LR2-5 cells were adsorbed to glow-discharged carbon-coated copper grids with Formvar films and imaged using a CM10 transmission electron microscope (Philips) coupled to a Gatan 792 BioScan camera. For details, see supplementary information.

### Phase contrast light microscopy, cell shape analysis and swimming analysis

*Hfx. gibbonsii* LR2-5 cells were imaged at 100× magnification using an Axio Observer.Z1 inverted microscope (Zeiss). Microscopy images were processed to analyze cell shapes using the FIJI/ImageJ plugin MicrobeJ.

Swimming analysis was performed at 63× magnification with an Axio Observer.Z1 inverted microscope (Zeiss). The movement of cells was recorded with 15 s time-lapse movies. For details, see supplementary information.

### Motility assay on semi-solid agar plates

Motility assays were performed as described previously (Quax *et al*., 2018; Li *et al*., 2020). For details, see supplementary information.

### Titration by spot on lawn assay and viral titer quantification

A dilution series of a virus preparation was prepared and 10 μL spots of each dilution were placed on the lawn of the known or other potential host cells *(Hfx. gibbonsii* LR2-5, *Haloferax gibbonsii* Ma2.38^T^ and *Hfx. volcanii* H26). Plates were incubated for 2-5 days at 37 °C and examined for the presence or absence of zones of growth inhibition. For details, see supplementary information.

The number of infectious viruses were determined by plaque assay by mixing 100 μL of virus dilutions with 300 μL dense host culture before plating in an overlay of MGM soft agar on MGM plates. Plates were incubated for 2-3 days at 37 °C. Plaques were counted and the number of infectious viruses per unit volume i.e. the titer (Plaque Forming Units/mL; PFU/mL) was determined.

### Identification of S-layer protein

A whole cell lysate of *Hfx. gibbonsii* LR2-5 was separated by SDS-polyacrylamide gel electrophoresis (SDS-PAGE) using a 7.5 % polyacrylamide-SDS gel. The gel was stained using Coomassie blue stain. A major band having the appropriate molecular weight was excised and used for identification by mass spectroscopy. For details, see supplementary information.

### Genome sequencing and assembly

Full details are provided in the supplementary information.

Cells of *Hfx. gibbonsii* LR2-5 were processed by Eurofins NGS Lab Constance (Constance) for DNA extraction. For PacBio RS sequencing, a “standard genomic library” was prepared and sequenced at Eurofins according to the manufacturer’s instructions. An automatic assembly using the HGAP3 pipeline was performed at Eurofins. To further improve the accuracy of the genome sequence, Illumina HiSeq sequencing was performed at Eurofins.

### Annotation of the *Hfx. gibbonsii* LR2-5 genome

Full details are provided in the supplementary information.

Gene prediction was performed using the RASTtk annotation server (Overbeek *et al*., 2014; Brettin *et al*., 2015; Lomsadze *et al*., 2018). The resulting annotation was curated using previously established procedures (Pfeiffer and Oesterhelt, 2015; Pfeiffer *et al*., 2020). The *Hfx. volcanii* annotation referred to as “up-to-date” is that from 6-JUN-2019, which is the basis for the community proteome project arcPP (Schulze *et al*., 2020).

An effort was made to reduce missing gene calls, especially small ones, by subjecting all intergenic regions −50 bp in the LR2-5 genome to an established BLASTx analysis procedure (Babski *et al*., 2016).

Annotation of stable RNAs and transposon analysis is described in the supplementary information.

### DNA methylation

Base modifications were analyzed using the SMRT® Analysis software version 7.0.1.66975 (Base Modification and Motif Analysis tool) (Chin *et al*., 2013). PacBio reads and the assembled genome sequence of strain LR2-5 were used as input. Results are is provided as supplementary information Table S4.

### Bioinformatic tools

Information on bioinformatic tools (e. g. MUMmer, BLAST, TYGS) can be found in the supplementary information.

## ACKNOWLEDGEMENTS

We thank Marina Geiger and Drishya S Gopan for the support with experiments. The TEM is operated by the University of Freiburg, Faculty of Biology, as a partner unit within the Microscopy and Image Analysis Platform, Freiburg.

## DATA AVAILABILITY STATEMENT

The nucleotide sequence accession numbers for the *Hfx. gibbonsii LR2*-5 genome are CP063205 (chromosome), CP063206 (plasmid pHGLR1), CP063207 (plasmid pHGLR2) and CP063208 (plasmid pHGLR3).

Raw reads have been deposited in the SRA archive under BioProject accession number PRJNA656928

## AUTHOR CONTRIBUTION

**Conceptualization**, T.E.F.Q, C.T, F.P.

**Data curation**, F.P.

**Project administration**, F.P., T.E.F.Q.

**Formal analysis**, F.P., M.D.S.

**Funding acquisition**, T.E.F.Q.

**Investigation**, **Validation**, S.S, F.P., M.R., C.T., H.M.O

**Supervision**, T.E.F.Q.

**Visualization**, S.S., C.T., M.R., M.D.S.

**Writing – original draft**, F.P., T.E.F.Q., C.T.

**Writing – review & editing**, F.P., T.E.F.Q., C.T., S.S., H.M.O., M.D.S.

