## Supplemental Information for "Cellular and genomic properties of *Haloferax gibbonsii* LR2-5, the host of euryarchaeal virus HFTV1"

### Supplemental Text

#### Strains of *Hfx. volcanii*

Two strains of Haloferax volcanii are relevant to the analysis, DS2$^T$ and H26. Strain DS2$^T$ is the wildtype strain of this species (Mullakhanbhai and Larsen, 1975). The laboratory strain H26 is an indirect offspring of DS2$^T$. First, DS2$^T$ was cured of the small plasmid pHV2, leading to strain DS70 (Wendoloski *et al.*, 2001). Then, the *pyrE2* gene was deleted from DS70, leading to strain H26 (Allers *et al.*, 2004). Thus, it is valid to use strain H26 for experimental analyses, e.g. studies on virus infectivity, but to compare the LR2-5 genome to that of the wildtype strain DS2$^T$.

#### Strand selection and setting of the point of ring opening for the LR2-5 plasmids.

The plasmid sequences were compared to the genomes from *Hfx. volcanii* DS2$^T$ and *Hfx. mediterranei* by BLASTn analysis. Long and highly conserved regions up to 16 kb with 95% DNA sequence identity were detected. Nevertheless, an unambiguous one to one correlation could not be discerned. Therefore, we compared the protein sequences of all Orc paralogs of *Hfx. volcanii* DS2$^T$ (*orc1* to *orc16*) to the *Hfx. gibbonsii* LR2-5 genome by tBLASTn analysis (see Table S3). This showed a close match of HVO_B0001 (Orc6) to plasmid pHGLR1 (100% protein sequence identity; HfgLR_20005, Orc2), a close match of HVO_A0001 (Orc3) to plasmid pHGLR2 (98% protein sequence identity; HfgLR_23005, Orc3), and a close match of HVO_C0001 (Orc10) to plasmid pHGLR3 (100% protein sequence identity; HfgLR_25005, Orc4). We next compared 1 kb of DS2$^T$ sequence upstream of these *orc* genes to the *Hfx. gibbonsii* LR2-5 genome by BLASTn. This revealed considerable conservation of the DNA sequence across the point of ring opening in the *Hfx. volcanii* DS2$^T$ plasmids. Conservation was close enough to allow the equivalent positioning of the points of ring opening for the plasmids from strain LR2-5.

#### Ribosomal RNA and tRNA genes.

The 16S rRNA shows 99 % sequence identity to that of the three analysed strains of *Hfx. gibbonsii* (strain ARA6: 4 point mutations; type strain Ma2.38, ATCC 33959, accession D13378: 6 point mutations, 3 one-base indels and *Hfx. volcanii* strain DS2$^{T}$: 10 point mutations). It should be noted that an alternative 16S rRNA sequence is available for *Hfx. gibbonsii* strain Ma2.38$^{T}$ via the draft genome sequence (AOLJ00000000), but that differs from the D13378 version as it lacks a few central bases, having an undefined base (N) at that position. This aberrant sequence precludes correct species assignment via the "16S rRNA based assignment" method in the TYGS server (see Methods).

There are tRNAs for all 20 amino acids. Three tRNA genes contain predicted introns (tRNA-Met, tRNA-Trp, tRNA-Gln). There is one aberrant complete tRNA (HfgLR_TRNA41) which differs from canonical tRNAs in the central region, so that an anticodon assignment is not possible. This aberrant tRNA occurs also in *Hfx. volcanii* (98 % nucleotide sequence identity) and *Hfx. gibbonsii* strain ARA6 (100 % nucleotide sequence identity). Curiously, the two largest plasmids (pHGLR1 and pHGLR2) each carry a single tRNA gene, specifying tRNA-iMet (HfgLR_TRNA65, HfgLR_TRNA66), and they are identical in sequence.

#### CRISPR-Cas loci

While *Hfx. volcanii* has a well-studied CRISPR-Cas system (Maier *et al.*, 2015, 2019), the three *Hfx. gibbonsii* strains differ in their carriage of this defence system. It is absent in strains LR2-5 and ARA6 but strain Ma2.38$^{T}$ carries a complete CRISPR-Cas system. These are organized as three CRISPR arrays, two of which have associated Cas proteins. There are 111 unique spacers (https://crisprcas.i2bc.paris-saclay.fr/CrisprCasFinder/Index). None of the CRISPR arrays contain any spacers specific to HFTV1 as determined by BLASTn. Curiously, one spacer in strain Ma2.38 targets the *hemQ* gene and the sequence is completely conserved in LR2-5, ARA6, and *Hfx. volcanii* DS2$^{T}$. Such self-targeting is tolerated in *Hfx. volcanii* (Stachler *et al.*, 2020). *Hfx. mediterranei* ATCC 33500 does have a spacer which closely matches a target in HFTV1. The target sequence is within HFTV1-gp26, the gene encoding the HK97 gp6-like/SPP1 gp15-like head-tail connector.

**Toxin Antitoxin Systems**

Toxin antitoxin systems (TA systems) are widely present in bacterial and archaeal genomes and consist of two protein components: toxins exert a toxic effect on the cell, while antitoxins can block this toxicity. Usually, antitoxins are metabolically more labile, such that the TA system can act as an 'addiction' module, when for example present on a plasmid (Gerdes *et al.*, 1986). TA systems are proposed to have a function in programmed cell death, stress response and abortive infection (Gerdes *et al.*, 2005; Tachdjian and Kelly, 2006). The latter is a phenomenon whereby cells commit altruistic suicide after viral infection in order to protect the clonal population (Pecota and Wood, 1996; Fineran *et al.*, 2009). The role of TA systems in archaea is less well characterized. They have been reported to be involved in stress response (Cooper *et al.*, 2009). Interestingly, infection by several crenarchaeal viruses lead to upregulation of the TA genes in *Sulfolbus* genomes (Ortmann *et al.*, 2008; Quax *et al.*, 2013; Erdmann *et al.*, 2014; León-Sobrino *et al.*, 2016).

In the LR2-5 genome, four TA loci are present, with two additional unlinked toxin genes and one additional antitoxin gene. Two of the TA loci belong to the VapB/C class and both unlinked toxins are predicted as VapC endonucleases, one of which is nonfunctional. The remaining two TA loci belong to the death-on-curing family (see Table S10). Most of the toxin and antitoxin genes are located on pHGLR1.

The nonfunctional VapC (HfgLR_12920) is the only toxin or antitoxin encoded on the chromosome. Interestingly, the *vapC* gene is located next to another nonfunctional gene which encodes an Abi/CAAX domain protein (HfgLR_12930). This indicates that these genes may be part of a no longer functional abortive infection system. Further downstream are an AbrB transcription regulator and a homolog to AbrB/VapB family protein (HfgLR_12950 and HfgLR_12955).

The HfgLR_20875/HfgLR_20880 death-on-curing/CHY (an uncharacterized protein annotated as "conserved hypothetical protein") TA system has a homolog (≥85 % protein sequence identity) in *Hfx. volcanii* (HVO_A0244/HVO_A0243); however, a closer homolog to the *Hfx. volcanii* antitoxin HVO_A0243 is also present in *Hfx. gibbonsii* LR2-5 as HfgLR_21655 (94 % protein sequence identity). This homolog does not have a predicted toxin in its vicinity.

One TA system (HfgLR_23165/HfgLR_23160) is encoded on pHGLR2. This system contains a VapC and a short uncharacterized protein. Both have very close homologs in both *Hfx. volcanii* (95 % and 93 % protein sequence identity) and in *Hfx. gibbonsii* ARA6 (100 % and 96 % protein sequence identity).

**Restriction Modification systems**

Analysis of the *Hfx. gibbonsii* LR2-5 genome showed that it lacks an ortholog of the Mrr family endonuclease HVO_0682 (*mrr*) from *Hfx. volcanii*. This restriction enzyme forms a barrier for foreign genetic elements that enter *Hfx. volcanii*. This is attributed to A methylation while C methylation does not trigger restriction (Holmes *et al.*, 1991). To obtain high transformation efficiency in *Hfx. volcanii*, genetic constructs are typically passaged though an *Escherichia coli dam* mutant to avoid A methylation and thus to overcome the restriction barrier. HVO_0682 shares a domain with *E. coli mrr* (37% protein sequence identity), which codes for a type IV restriction enzyme which acts on DNA methylated at A or C residues (Waite-Rees *et al.*, 1991). Deletion of HVO_0682 allows efficient transformation with methylated DNA, confirming the predicted function of the enzyme (Allers *et al.*, 2010). The majority of haloarchaea, including *Hfx. gibbonsii* ARA6, lack orthologs to HVO_0682 and its presence in *Hfx. volcanii* seems to be an exception.

*Hfx. volcanii* possesses another gene (HVO_0803) with an Interpro-assigned *mrr* domain but this does not hit *E. coli mrr* upon BLASTp analysis. Because HVO_0803 has a centrally located transmembrane domain, it may not be involved in restriction. The LR2-5 ortholog of HVO_0803 is HfgLR_04065.

In a previous survey, a patchy distribution of restriction/modification systems in haloarchaea was observed and it was concluded that they are not a main countermeasure against virus infection (Fullmer *et al.*, 2019). Consistent with this, type I restriction/modification systems are found in both, *Hfx. volcanii* DS2$^T$ and *Hfx. gibbonsii* LR2-5, but these are only marginally related to each other. One of the LR2-5 type I restriction/modification systems is encoded on IGE1 (see above). There is one potential DNA modification methylase in strain LR2-5 and three in *Hfx. volcanii*, again with only marginal similarities (Table S11). Furthermore, as described above, four methylated sequence motifs are present in *Hfx. gibbonsii* LR2-5 (Table S4) but the responsible DNA modification methylases have not yet been unraveled. Strain LR2-5 does not have additional proteins with restriction endonuclease domains while *Hfx. volcanii* has two such proteins which, however, have not yet been experimentally characterized.

### Supplemental Materials and Methods

#### Media and growth conditions

*Haloferax gibbonsii* LR2-5 cells were cultured aerobically at 37 °C, 42 °C, or 45 °C under constant rotation at 120 rpm in medium prepared with 30 % (vol/vol) artificial salt water (SW) containing (per liter) 240 g of NaCl, 30 g of $MgSO_4 \cdot 7 H_2O$, 35 g of $MgCl_2 \cdot 6 H_2O$, 7 g of KCl, 80 mM Tris HCl (pH 7.2) and 5 mM $CaCl_2$. The 30 % stock of SW was diluted to make the working media with either 18 % (vol/vol), 20 % (vol/vol) or 23 % (vol/vol) SW. For growth in rich YPC medium (Yeast extract, Peptone, Casamino acids) (Allers *et al.*, 2004) the salt water was supplemented with 0.5 % (wt/vol) yeast extract (Difco), 0.1 % (wt/vol) peptone (Oxoid), and 0.1 % (wt/vol) casamino acids (Difco). For growth in rich MGM (modified growth medium) (Nuttall and Smith, 1993) yeast extract 0.5 %, (wt/vol) and peptone 0.1 % (wt/vol) were added. For growth in selective Casamino Acids medium (CA) (Allers *et al.*, 2004) casamino acids were added to the SW preparation to a final concentration of 0.5 % (wt/vol). Additionally, thiamine and biotin were added to final concentrations of 0.9 mg/L and 0.1 mg/L, respectively. CA medium modified with trace element solution (CAB) (Duggin *et al.*, 2015) was prepared in the same manner as CA medium with addition of 1 % (vol/vol) expanded trace element solution. The trace element solution contains (per 100 mL) 36 mg $MnCl_2 \cdot 4 H_2O$, 44 mg $ZnSO_4 \cdot 7 H_2O$, 230 mg $FeSO_4 \cdot 7 H_2O$, 5 mg $CuSO_4 \cdot 5 H_2O$ (filter sterilized). The medium was autoclaved (pH 7.2 adjusted with KOH). After cooling, $CaCl_2$ was added to a final concentration of 3 mM. For recording of growth curves, the growth was started by diluting the culture to an $OD_{600}$ of 0.001. The experiments were performed in triplicate (containing 3 biological replicates) for each medium, temperature and salt condition.

#### Transmission electron microscopy

*Haloferax gibbonsii* LR2-5 cells were grown overnight at 37 °C in CA medium (18% SW) to an $OD_{600}$ of ~0.1 and harvested by centrifugation at 2,000 g for 15 min. The cells were concentrated and resuspended in CA medium before being adsorbed to glow-discharged carbon-coated copper grids with Formvar films for 10 s. The samples were washed three times in drops of sterile 2 M NaCl and subsequently stained for 15 seconds with 2 % (wt/vol) uranyl acetate. The uranyl acetate solutions were prepared in 2 M NaCl and sterile filtered. Cells were imaged using a CM10 transmission electron microscope (Philips) coupled to a Gatan 792 BioScan camera.

#### Phase contrast light microscopy, cell shape analysis and swimming analysis

*Haloferax gibbonsii* LR2-5 cells were grown in CA medium containing 18% SW at 42 °C. For imaging, cultures were diluted to an $OD_{600}$ of 0.1 and 5 µL cell suspension was spotted on an agarose pad (0.4 % (wt/vol) agar, 18 % SW) and covered with a glass slip. For shape analysis, cells were imaged at 100× magnification in the Phase contrast mode (PH3) using an Axio Observer.Z1 inverted microscope (Zeiss). Microscopy images were processed to analyze cell shapes using the FIJI/ImageJ plugin MicrobeJ.

For swimming analysis, one milliliter of culture diluted to an $OD_{600}$ of 0.1 was pipetted into a round DF 0.17-mm microscopy dish (Bioptechs) and observed at 63× magnification in the DICII mode with an Axio Observer.Z1 inverted microscope (Zeiss) equipped with a heated (45 °C) XL-5 2000 incubator running VisiVIEW software. The movement of cells was recorded with 15 s time-lapse movies.

#### Motility assay on semi-solid agar plates

Motility assays were performed as described previously (Quax *et al.*, 2018; Li *et al.*, 2020). Semi-solid agar plates were made in YPC, MGM, CA, and CAB medium containing 0.3 % (wt/vol) agar and either 18 % or 23 % SW. Fresh cells from agar plates were grown in 18 % CA medium. The cultures were diluted to an $OD_{600}$ of 0.5 and about 10 µL of the inoculum was spotted on a semi-solid agar plate. The experiments were performed in at least triplicate (containing 3 biological replicates) per condition. The motility ring formed after cultivation of plates at 45 °C for 3 days was analyzed by scanning of plates and measurement of the diameter using FIJI/ImageJ.

#### Preparation of virus stock

For preparation of virus stock, semi-confluent plates were produced using the double-layer method (Mizuno *et al.*, 2019). Agar (Bacto-Agar) was added to prepare solid (14 g/L) or top-layer (4 g/L) media containing 20 % and 18 % (wt/vol) SW, respectively. Plates were incubated for 2 nights at 37 °C after which the top-layer agar was collected and incubated in MGM medium (2 mL per plate) at 37 °C for 1.5 h with shaking. Viral particles were harvested by centrifugation of the agar (8,000 g, 20 min, 4 °C). The supernatant was stored in the dark at 4 °C.

#### Titration by spot on lawn assay

*Haloferax gibbonsii* strain Ma2.38$^{T}$ and *Hfx. gibbonsii* strain LR2-5 were grown aerobically at 37 °C in MGM. Agar (Bacto-Agar) was added to prepare solid (14 g/L) or top-layer (4 g/L) media containing 20 % and 18 % (wt/vol) SW, respectively. 300 µL dense host culture were mixed with 3 mL of melted soft agar and mixed thoroughly before being poured on a MGM plate. The lawn was solidified and dried at room temperature for an hour. A dilution series of a virus preparation was prepared and 10 µL spots of each dilution were placed on the host lawn. MGM was used as a control. Plates were incubated for 2-5 days and examined for the presence or absence of zones of growth inhibition. The experiment was performed on parallel plates and was repeated for at least three times.

#### Viral titer quantification

Plaque assays were performed to calculate the viral titer. 100 µL of virus dilutions were mixed with 300 µL dense host culture before plating in an overlay of MGM media and agar. Plates were incubated for 2-3 days at 37 °C. Plaques were enumerated and the number of infectious viruses i.e. the titer (Plaque Forming Units/mL) was determined.

#### Identification of S-layer protein

A whole cell lysate of *Hfx. gibbonsii* strain LR2-5 was obtained by pelleting 1 mL of culture and resuspending in 80 µL dd$H_2O$, 20 µL 5 x SDS loading dye and heating to 95 °C for 20 min. 30 µL lysate were loaded onto a 7.5 % polyacrylamide-SDS gel which was run at 150V until the front exited the gel. The gel was stained using Coomassie blue stain.

A band at the appropriate position was excised and cut into small (1 $mm^3$) pieces. The gel piece was then treated with 10 mM $NH_4HCO_3$ for 10 min at room temperature. The supernatant was removed, the sample was treated again with 5 mM $NH_4HCO_3$ in 60 % (vol/vol) ethanol for 10 min at room temperature. These steps were repeated 3 times. The sample was then treated with 10 mM DTT (dithiothreitol) in 10 mM $NH_4HCO_3$ for 30 min at 56 °C. The supernatant was removed and the sample was incubated with 50 mM IAA (iodoacetamide) in 10 mM $NH_4HCO_3$ for 30 min at RT in the dark. The supernatant was removed, the sample was incubated with 10 mM $NH_4HCO_3$ for 10 min at RT. The supernatant was removed, the sample was incubated with 100 % ethanol for 10 min at RT. The steps after IAA treatment were repeated twice. Gel pieces were dried via speedvac (35 °C, 1h). Trypsin (0.6 µg) was added to the sample and incubated at 37 °C overnight. 0.05 % TFA (trifluoroacetic acid) in 50 % ACN (acetonitrile) were added, the samples were sonicated on ice for 10 min. The supernatant was transferred to a new tube. The steps after trypsin digestion were repeated, the supernatants were combined. The supernatants were dried via speedvac (35 °C, 1.5 h). Samples were resuspended in 0,1% TFA (in $H_2O$) and sonicated at RT for 3 min. Samples were centrifuged at 12,000 RPM for 5 min, the supernatant was transferred to a fresh UPLC tube and subjected to LC/MS measurement.

#### Genome sequencing and assembly

Cells of *Hfx. gibbonsii* LR2-5 were grown aerobically at 45ºC under shaking (120 rpm) to an $OD_{600}$ of 1.0 in YPC medium (Allers *et al.*, 2004). Cells collected by centrifugation for 30 min at 5,000 g, room temperature were processed by Eurofins NGS Lab Constance (Constance) for DNA extraction. The cells were resuspended in ST buffer (1 M NaCl, 20 mM Tris-HCl, pH 7.5). Next, lysis solution (100 mM EDTA pH 8.0, 0.2 % SDS) was added to the cells, and 1 mL ethanol was carefully layered on top of the aqueous solution, generating two phases. DNA was spooled at the interface onto a capillary. The DNA was washed in ethanol and resuspended in 500 µl TE buffer (10 mM Tris-HCl pH 8.0, 10 mM EDTA). DNA was precipitated by adding 50 µl 3 M sodium acetate (pH 5.2) and 400 µL isopropanol and collected by centrifugation (16,000 g, 5 min). The pellet was washed with 70% ethanol and dried and the DNA was resuspended in 100 µl TE buffer. The sample was treated with RNAse A (1 µL of 30 mg/mL in 50 % glycerol, Sigma-Aldrich Cat. R 4642: 45°C for ≥1 hour). DNA was left at 4°C overnight to resuspend completely.

For PacBio RS sequencing, a "standard genomic library" was prepared at Eurofins according to the manufacturer's instructions. The data originated from 1 SMRT cell. We obtained 99,807 reads totaling 846,204,000 bp with a mean read length of 8,478 bp (N50: 12,269 bp).

An automatic assembly using the HGAP3 pipeline was performed at Eurofins, based on 87,906 reads totaling 774,559,018 bp with a mean length of 8,811 bp (N50: 12,661 bp). The assembly resulted in four contigs, which showed terminal redundancy and thus represented distinct circular replicons (1 chromosome and 3 large plasmids). The mean coverage was 150-fold for the chromosome and the largest plasmid pHGLR1, 120-fold for the 2$^{nd}$ plasmid pHGLR2, and 290-fold for the 3$^{rd}$ (smallest) plasmid pHGLR3.

To further improve the accuracy of the genome sequence, Illumina HiSeq sequencing was performed and produced 2.42 Gb of raw sequence consisting of 8,073,870 reads in paired-end mode. A total of 2.505 million high quality paired-reads (read length=150 bp; coverage=94-fold) were selected (from the ~8 million reads returned) and mapped to the four contigs obtained from PacBio sequencing using the 'map to reference' option and Geneious mapper tool (default settings) in the Geneious (version 10.2.6 http://www.geneious.com/). Alignments were examined manually for discrepant base calls and corrections made where required.

##### Assignment of strain LR2-5 to the species *Hfx. gibbonsii*

The whole genome sequence of strain LR2-5 was submitted to the Type Strain Genome Server (TYGS; https://tygs.dsmz.de) (Meier-Kolthoff and Göker, 2019), which assigned the strain to the *Haloferax gibbonsii* species (see also main text). However, there is lack of confirmation by the "16S rRNA based assignment" method in the current version of the TYGS server, which could be attributed to a 16S rRNA sequence problem for the type strain of *Hfx. gibbonsii*. This strain (Ma2.38; ATCC 33959) is represented by a draft genome sequence (accession AOLJ01000000, 30 contigs) (Becker *et al.*, 2014). The 16S rRNA in the draft genome deviates from that previously obtained for ATCC 33959 (accession D13378), including an internal deletion associated with an undefined base (N), which indicates reduced reliability (see also below).

##### Annotation of protein-coding genes of the *Hfx. gibbonsii* LR2-5 genome

Gene prediction was performed using the RASTtk annotation server (Overbeek *et al.*, 2014; Brettin *et al.*, 2015; Lomsadze *et al.*, 2018). The resulting annotation was curated against an up-to-date Gold Standard Protein based annotation of the *Hfx. volcanii* genome, as well as the annotation of more than 10 additional haloarchaeal genomes (Pfeiffer and Oesterhelt, 2015). Further annotation enhancements had been made when annotating the *Halobacterium salinarum* type strain (strain 91-R6) genome (Pfeiffer *et al.*, 2020). The annotation referred to as "up-to-date" is that from 6-JUN-2019, which is the basis for the community proteome project arcPP (Schulze *et al.*, 2020)

For manual curation, each protein-coding gene was compared with BLASTp to the set of Gold Standard Protein annotated haloarchaeal genomes, which was supplemented with the UniProt proteome from *Hfx. gibbonsii* strain ARA6. Typically, besides the ortholog from strain ARA6, the best match was from *Hfx. volcanii* with an average of 93% protein sequence identity. Any start codon assignment discrepancies were immediately obvious from the alignment and were resolved. All genes with a TTG start codon were evaluated. Also, disrupted genes were detected and resolved during this analysis. The annotation of the *Hfx. volcanii* ortholog was copied to the corresponding protein from strain LR2-5. If the *Hfx. volcanii* protein had the lowest annotation level (conserved hypothetical protein; called "uncharacterized protein" in GenBank und UniProt), the UniProt entry was inspected for the assignment of InterPro domains and the protein name was improved if appropriate.

All proteins specific for the LR2-5 genome were annotated by comparison to (a) the set of carefully annotated haloarchaeal genomes (see above), (b) the SwissProt section of UniProt and (c) the *Hfx. gibbonsii* strain ARA6 proteome, including associated InterPro domains.

An effort was made to reduce missing gene calls, especially of small proteins, by subjecting all intergenic regions ≥50 bp in the LR2-5 genome to a BLASTx analysis procedure (Babski *et al.*, 2016).

#### Annotation of stable RNAs

Initially, the annotation of stable RNAs (rRNAs, tRNAs, RNase P RNA, 7S RNA, H/ACA guide RNA) from *Hfx. volcanii* was made consistent with RFAM (Kalvari *et al.*, 2018), following a procedure previously described for *Halobacterium* (Pfeiffer *et al.*, 2020). Stable RNAs from strain LR2-5 were then annotated using BLASTn comparison to the stable RNAs from *Hfx. volcanii*. For tRNAs, the results were found to be consistent with results from the tRNAscan-SE server (http://lowelab.uscs.edu/tRNAscan-SE) (Lowe and Eddy, 1997; Chan and Lowe, 2019).

##### Transposon analysis

Transposons were identified by BLASTn and BLASTx comparison to an extensive in-house collection of haloarchaeal transposons, and to the ISFinder database (Siguier *et al.*, 2006, 2012) by a previously described procedure (Pfeiffer *et al.*, 2020). Identified transposons were added to the in-house database and were used for a subsequent iterative transposon analysis using BLAST. Newly identified transposons were submitted to and accepted by ISFinder.

#### DNA methylation

Base modifications were analyzed using the SMRT® Analysis software version 7.0.1.66975 (Base Modification and Motif Analysis tool) (Chin *et al.*, 2013). PacBio reads and the assembled genome sequence of strain LR2-5 were used as input. For results see Table S4.

#### Additional bioinformatics tools

As general tools, MUMMER (Delcher *et al.*, 2003) and the BLAST suite of programs (Altschul *et al.*, 1997; Johnson *et al.*, 2008) were used for genome comparisons. The CRISPR finder web server (http://crispr.i2bc.paris-saclay.fr) was used to analyze for CRISPR elements in strains of *Hfx. gibbonsii* (Grissa *et al.*, 2008). Prophage searches were performed using PHASTER (http://phaster.ca/) and Profinder (http://aclame.ulb.ac.be/Tools/Prophinder/) but none were identified. Circular genome maps were created using the CGView Server (http://stothard.afns.ualberta.ca/). Genomic island (GI) prediction used Island Viewer 4 (http://www.pathogenomics.sfu.ca/islandviewer/) described by (Bertelli *et al.*, 2017).

Glycosylation sites were predicted using https://www.hiv.lanl.gov/cgi-bin/GLYCOSITE/glycosite_main.cgi (Zhang *et al.*, 2004). Tetranucleotide frequency analysis was performed using https://www.cmbl.uga.edu//software/signature.html. Type III signal sequences were detected by FlaFind (version 1.2; http://signalfind.org/flafind.html; (Szabó *et al.*, 2007))

### Supplemental Figures

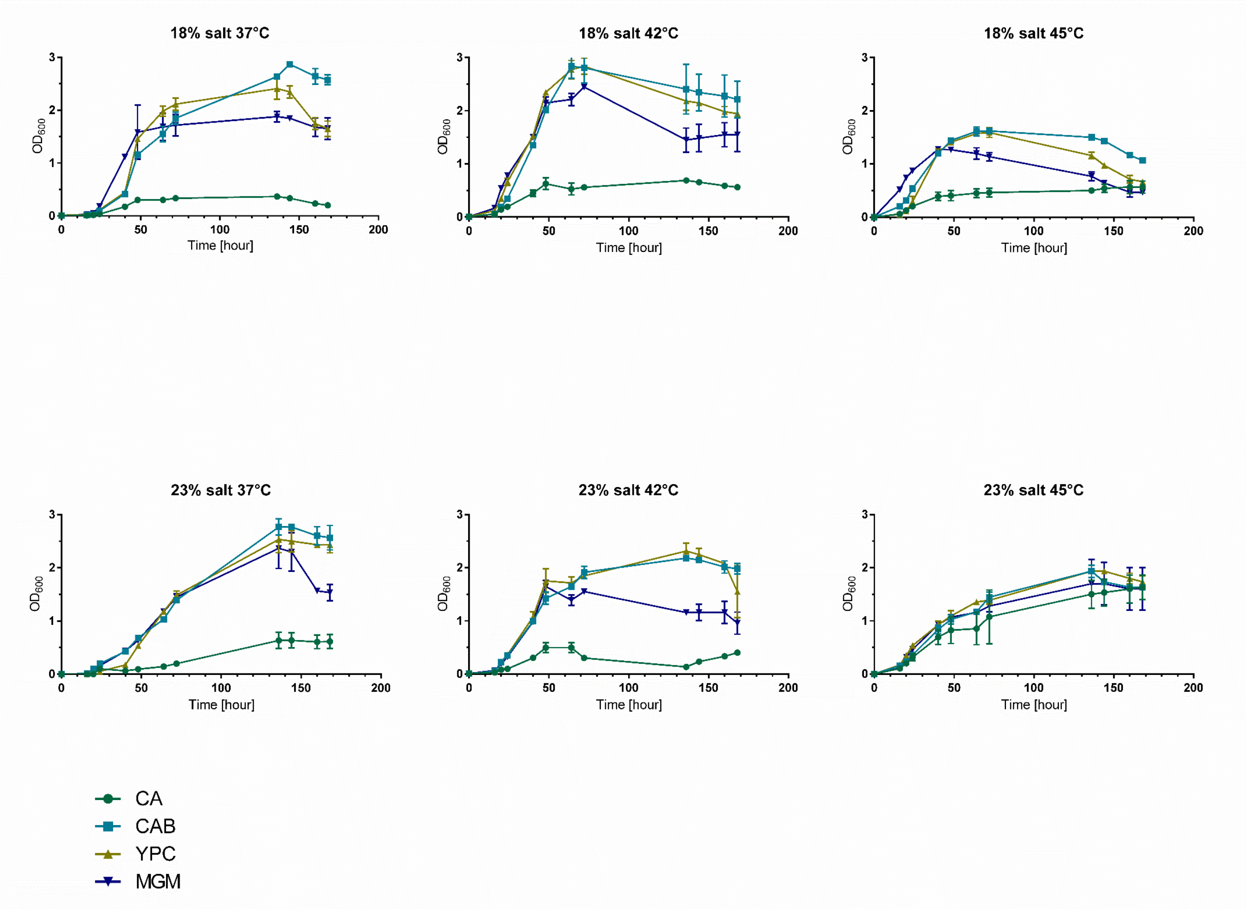


**Figure S1**. **Growth analysis of *Hfx. gibbonsii* LR2-5 in different media and temperatures.**

Growth curves of *Hfx. gibbonsii* LR2-5 were reported over a period of 160 hours. Cells were grown in 18 % or 23 % SW (wt/vol) salt concentration. Shown are the growth curves in selective media: CA (green circles), CAB (light blue rectangles) and in rich media: YPC (yellow triangles) and MGM (blue inverted triangles). Growth yields are dependent on the temperature and nutrient composition of the media. *Haloferax gibbonsii* LR2-5 grows best in YPC medium and CA supplemented with trace elements. Cells grown at 37°C exhibit prolonged lag-phases compared to cells grown at higher temperatures. Lag phase at 42 °C resemble that at 45 °C. We found a growth optimum at 42 – 45 °C. Cultures grown at 45°C reach lower final optical densities. Average optical density at 600 nm ($OD_{600}$) was calculated from three independent technical replicates. Each figure represents one out of three biological replicates with similar results. Error bars represent standard deviation from three technical replicates.

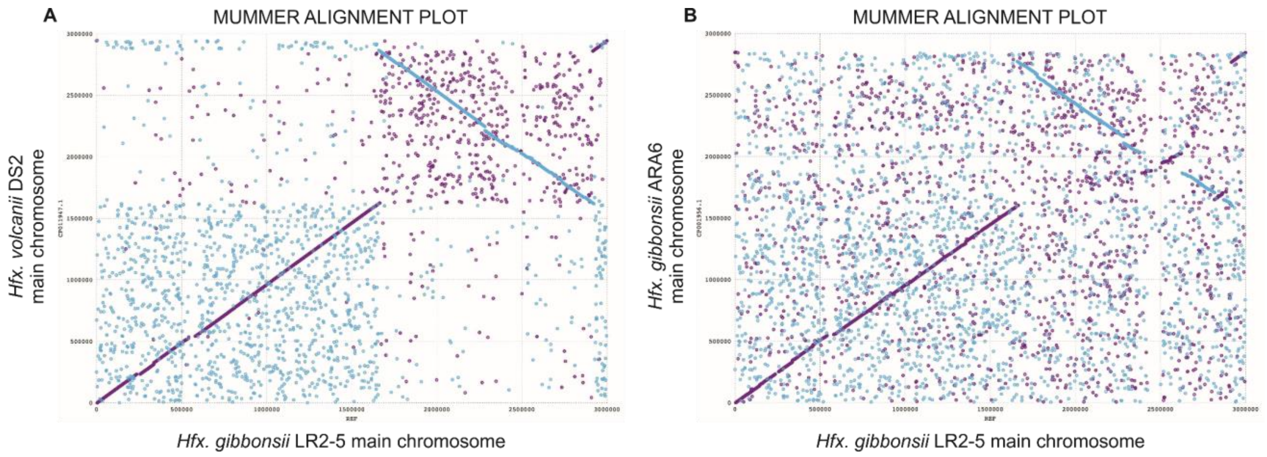


**Figure S2**. **MUMmer comparison of *Hfx. gibbonsii* LR2-5 chromosome to those of closely related genomes.** Dotplot comparisons (MUMmer 3.23) of the LR2-5 chromosome sequence with the chromosomes of (A) *Hfx. volcanii* DS2$^{T}$ and (B) *Hfx. gibbonsii* ARA6. Purple and blue diagonal lines indicate nucleotide sequence similarity; purple indicates colinear matches and blue indicates reverse matches. The inversion, which is flanked by the two rRNA operons, can be attributed to LR2-5 as the gene order corresponds between ARA6 and DS2$^{T}$.

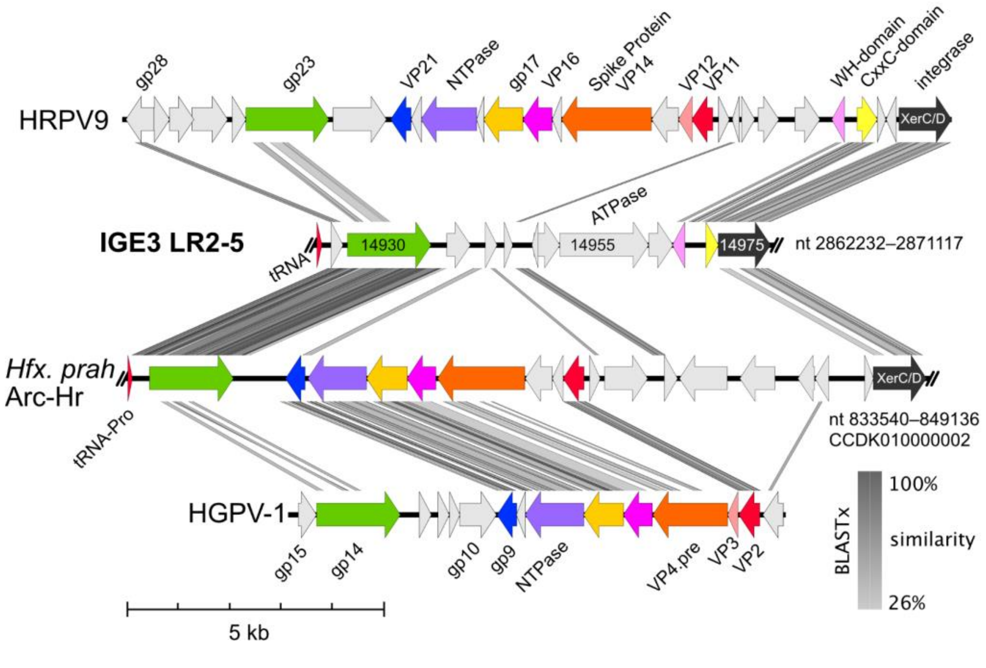


**Figure S3. Comparison of the IGE3 region of strain LR2-5 with haloviruses HRPV9 and HGPV-1 and with a related integrated element in *Hfx. prahovense* strain Arc-Hr.** Similar protein coding regions are the same colour across the different gene maps. BLASTx similarities of the corresponding encoded proteins are shown by grey shading (similarity scale shown at lower right). Halorubrum pleomorphic virus 9 (HRPV9; accession KY965934). Halogeometricum pleomorphic virus 1 (HGPV-1; accession JN882267). Halovirus annotations are shown above (HRPV9) or below (HGPV-1) their respective maps. IGE3 LR2-5, integrative genetic element 3 of *Hfx. gibbonsii* strain LR2-5. Nucleotide range given at right end. Locus tag numbers shown inside gene arrows, e.g. 14930 = HfgLR_14930. *Hfx. prah* Arc-Hr, *Haloferax prahovense* strain Arc-Hr (accession LK053000, contig CCDK010000002; previously called *Hfx. alexandrinus*). Annotations were downloaded from IMG/JGI (IMG ID Ga0056861_102). Nucleotide range given at right end. Scale bar is shown below the diagram.

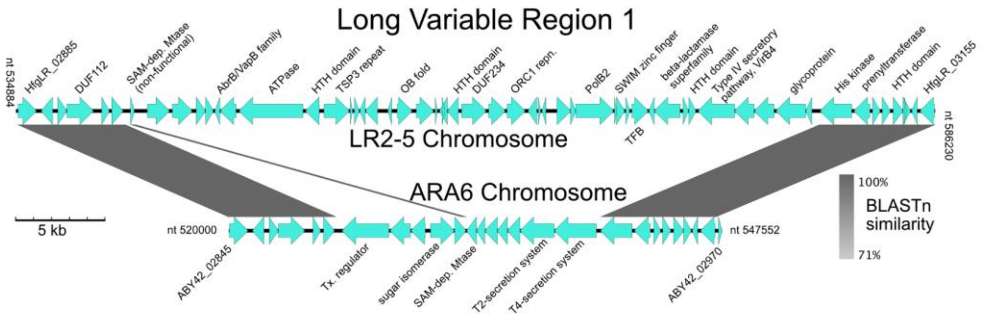


**Figure S4. Comparison between LR2-5 and ARA6 genomes across Long Variable Region 1 (LVR1).** Between the flanking regions that show high nucleotide similarity (BLASTn, E-value $\leq 10^{-20}$) are two regions showing little similarity. In this region, the two genomes differ in length and gene composition. From the truncated SAM-dependent methyltransferase gene (SAM-dep. Mtase) in LR2-5, it might be inferred that LR2-5 has suffered a deletion of the five genes found in ARA6. The other genes differ in sequence and number and appear to have been the result of a replacement event.

#### Supplemental Tables

**Table S1. Swimming of LR2-5 at different ODs.** Motility level indicators: -: 1-4 %, +/-: 5-29 %, +: 30-49 %, ++: 50-69 %, +++: 70-100 % swimming cells. N.A: not applicable. Data obtained via microscopy-based swimming analysis performed at 45°C. Media names and their compositions are given in Materials and Methods.

| | **Optical Density at 600 nm ($OD_{600}$)** | | | | |
|---|---|---|---|---|---|
| **Medium** | **0.01 - 0.1** | **0.1 - 0.3** | **0.3 - 0.7** | **0.7 - 1.2** | **1.2 - 1.8** |
| **CA** | +++ (80%) | ++ (60%) | ++ (30%) | + (5%) | N/A |
| **CAB** | +++ (85%) | +++ (80%) | ++ (60%) | + (30%) | +/- (5%) |
| **YPC** | + (25%) | + (10%) | +/- (5%) | - (1%) | - (1%) |
| **MGM** | +++ (70%) | ++ (60%) | ++ (40%) | ++ (40%) | +/- (5%) |

**Table S2. Replicons of *Hfx. gibbonsii* LR2-5.** Core statistical data and summary are given for the replicons. For proteins, the total number is given and how many of these are disrupted (pseudogenes). Protein-coding genes which have been targeted by a transposon (or other mobile genetic element) are annotated (and thus counted) as multiple single-region ORFs. The ordered locus tag for the first and last regular protein-coding gene of each replicon is indicated. Only the digit part is given, the prefix (HfgLR) is omitted. Ordered locus tags below 20,000 are used for the chromosome. For each plasmid, the serial number of the 1$^{st}$ gene ends with 005. Each rRNA operon has 3 rRNAs (16S, 23S, 5S). ncRNAs are the 7S RNA, RNAse P RNA and the H/ACA guide RNA. Regular tRNAs are those suitable for translation. There is one atypical tRNA without anticodon assignment and several partial tRNAs. Average read coverage is given for PacBio long reads and for a subset (2.5 million out of 8.0 million) of Illumina reads. The relative coverage compared to that of the chromosome is indicated in parenthesis.

| | **replicons** | | | | **summary** | |
|---|---|---|---|---|---|---|
| replicon | chromosome | pHGLR1 | pHGLR2 | pHGLR3 | plasmids | genome |
| length (bp) | 2,999,641 | 608,598 | 322,970 | 65,035 | 996,603 | 3,996,244 |
| GC (%) | 66.9 | 61.7 | 66.9 | 55.8 | 63.0 | 66.0 |
| proteins (total) | 3116 | 579 | 301 | 68 | 948 | 4064 |
| pseudogenes (total) | 27 | 35 | 2 | 11 | 48 | 75 |
| locus tag ranges | 00005-15580 | 20005-22895 | 23005-24505 | 25005-25340 | - | - |
| rRNA operons (total) | 2 | 0 | 0 | 0 | 0 | 2 |
| ncRNAs (total) | 3 | 0 | 0 | 0 | 0 | 3 |
| regular tRNAs (total) | 53 | 1 | 1 | 0 | 2 | 55 |
| fragment/aberrant tRNA (total) | 10 | 1 | 0 | 1 | 2 | 12 |
| average read coverage (PacBio) | 149 (1.00) | 150 (1.01) | 117 (0.79) | 290 (1.95) | - | - |
| average read coverage (Illumina) | 93 (1.00) | 94 (1.01) | 75(0.81) | 221(2.38) | - | - |

**Table S3. The Orc1-type DNA replication proteins of *Hfx. gibbonsii* LR2-5, *Hfx. volcanii* DS2$^T$ and *Hfx. gibbonsii* ARA6.** For LR2-5 and DS2$^T$, gene names and locus tags are included and the listing is complete. Orthologs are listed in the same line, including the level of protein sequence identity (seqid). The absence of an ortholog is indicated by a dash. One distant ortholog is highlighted in yellow. For *Hfx. gibbonsii* ARA6, only the locus tag and protein sequence identity to strain LR2-5 are given and no attempt was made to identify *orc* genes that lack orthologs in strains LR2-5 and DS2$^T$. For strain LR2-5, the first four gene serials were assigned to the *orc* genes at the start of the replicons (chromosome and the three plasmids). The remainder of the serials was assigned along the genome.

| ***Hfx. gibbonsii* LR2-5** | | ***Hfx. volcanii* DS2$^T$** | | | ***Hfx. gibbonsii* ARA6** | |
|---|---|---|---|---|---|---|
| **gene** | **locus tag** | **gene** | **locus tag** | **seqid(%)** | **locus tag** | **seqid(%)** |
| *orc1* | HfgLR_00020 | *orc1* | HVO_0001 | 94 | ABY42_00005 | 99 |
| *orc2* | HfgLR_20005 | *orc6* | HVO_B0001 | 100 | ABY42_15465 | 100 |
| *orc3* | HfgLR_23005 | *orc3* | HVO_A0001 | 98 | ABY42_18045 | 100 |
| *orc4* | HfgLR_25005 | *orc10* | HVO_C0001 | 100 | - | - |
| *orc5* | HfgLR_00955 | *orc9* | HVO_0194 | 99 | ABY42_00915 | 99 |
| *orc6* | HfgLR_03020 | *orc12* | HVO_A0072 | 52 | - | |
| *orc7* | HfgLR_03220 | *orc2* | HVO_0634 | 99 | ABY42_03020 | 100 |
| *orc8* | HfgLR_07785 | *orc15* | HVO_1537 | 90 | ABY42_07470 | 99 |
| *orc9* | HfgLR_08770 | *orc5* | HVO_1725 | 98 | ABY42_08435 | 100 |
| *orc10* | HfgLR_12880 | - | - | - | - | - |
| *orc11* | HfgLR_13010 | - | - | - | - | - |
| *orc12* | HfgLR_13465 | *orc16* | HVO_2133 | 92 | ABY42_10115 | 99 |
| *orc13* | HfgLR_21430 | *orc7* | HVO_A0257 | 99 | - | - |
| *orc14* | HfgLR_21550 | *orc8* | HVO_C0057 | 99 | ABY42_18895 | 99 |
| *orc15* | HfgLR_21930 | - | - | - | ABY42_18725 | 98 |
| *orc16* | HfgLR_21995 | - | - | - | - | - |
| - | - | *orc4* | HVO_2042 | - | - | - |
| - | - | *orc14* | HVO_2292 | - | - | - |
| - | - | *orc11* | HVO_2293 | - | - | - |
| - | - | *orc13* | HVO_A0064 | - | - | - |

**Table S4. DNA modifications of *Hfx. gibbonsii* LR2-5.** DNA modifications were detected in PacBio reads using the SMRT® Analysis software (see Methods). Modified bases are in bold, with the modification type indicated in superscript. Rows 3 and 4, and rows 5 and 6 represent pairs of complementary sequences where both are written in the 5' to 3' direction. Ranges of % site methylation show differences in estimates across the four replicons.

| Motif and modification | Modified strand | sites methylated (%) | Motifs in genome (total) |
|---|---|---|---|
| $^{m4}$**C**TAG | both | 76-86 | 932 |
| GCG$^{m4}$**C**TG | one | 95-98 | 4399 |
| $^{m6}$**A**YCnnnnnCTTYG<br>CRA$^{m6}$**A**GnnnnnGRT | both | 100 | 849 |
| CC$^{m6}$**A**nnnnnnRTCnC<br>GnG$^{m6}$**A**YnnnnnnTGG | both | 100 | 683 |

**Table S5. Integrative Genetic Elements (IGE) and Long Variable Regions (LVR) of strain LR2-5.** For each IGE and LVR, the position is shown with the covered proteins indicated in parenthesis (only the digit part of the ordered locus tag is given, the prefix HfgLR is omitted). For IGEs, an orientation is indicated, based on the strand encoding the XerC/D integrase homolog, which always was found close to 3`end of the associated complete tRNA. The locus tag for the tRNA is indicated, omitting the prefix HfgLR. The partial tRNA at the other end of IGE2 is annotated as HfgLR_TNA39, the other partial tRNA duplications were too short to become annotated.

| **Name** | **Length (bp)** | **Position (nt)** <br> **(protein locus tags)** | **Integrase** | **tRNA** <br> **(tRNA locus tag)** | **Terminal direct repeat (bp)** |
|---|---|---|---|---|---|
| IGE1 | 20,300 | 236046-256345/F (01305-01385) | XerC/D | tRNA-Pro (TRNA03) | 18 |
| IGE2 | 11,508 | 1776563-1765056/R (09310-09350) | XerC/D | tRNA-Glu (TRNA40) | 42 |
| IGE3 | 8,886 | 2862232-2871117/F (14925-14975) | XerC/D | tRNA-Arg (TRNA61) | 18 |
| LVR1 | 38,047 | 541301-579348 (02915-03115) | - | - | - |
| LVR2 | 194,625 | 2349993-2544618 (12450-13295) | - | - | - |

**Table S6. Mobile genetic elements (MGE) of *Hfx. gibbonsii* LR2-5 and ARA6.** Only complete MGEs are listed. Complete MGEs have both termini intact and lack long internal deletions. For each element, a **class** is assigned (ISH, MITE, or solo), a **type**, a **name**, and the number of copies in each strain (#) is given. The existence of multiple near-identical copies, indicative of transposition (mobilization), is a hallmark of MGEs. Such expanded elements are absent from both strains of *Hfx. gibbonsii*. Also, most MGEs are strain-specific, and only few occur in both strains. MGEs of class ISH carry a transposase gene, which determines the assignment of the transposon type. MGEs of class MITE are non-coding and are mobilized by a transposon of the same type in trans. The assignment of the MITE type is based on the sequence of the inverted terminal repeat. Transposons and MITEs of type IS1341 (and also IS605 and IS607) lack inverted terminal repeats but instead have near-terminal palindromes at both ends. IS1341 class solo are short sequences, typically ca 52 bp long, and consist of a single copy of the palindromic sequence. Transposon and MITE names follow ISFinder rules. Complete transposons with an intact transposase gene, which are assigned to *Hfx. gibbonsii*, are named ISHgi with a serial number. Transposons submitted to and accepted by ISFinder are in bold. Those submitted to ISFinder but not yet accepted are underlined. According to ISFinder rules, transposons with >95% nucleotide sequence identity are the same element, even if they occur in other species (e.g. ISHbo1, ISHvo18; highlighted in grey). Transposons named HgiIRS (Insertion-Related Sequence) with a serial number are not suitable for ISFinder submission because the transposase gene is disrupted. Equivalent rules apply to MITEs named MITEHgi and e.g. MITEHvo. MGEs of class solo are not suitable for ISFinder submission. Also, elements of type NmIRS44 are not well enough defined to be suitable for ISFinder submission.

| | | LR2-5 | | ARA6 | |
|---|---|---|---|---|---|
| **type** | **class** | **name** | **#** | **name** | **#** |
| ISH9 | ISH | **ISHgi4** | 2 | - | 0 |
| ISH9 | ISH | **ISHgi5** | 1 | - | 0 |
| ISH9 | ISH | HgiIRS40 | 1 | - | 0 |
| ISH9 | MITE | - | 0 | MITEHvo1 | 1 |
| ISH9 | MITE | - | 0 | MITEHvo3 | 2 |
| ISH9 | MITE | - | 0 | **MITEHgi1** | 1 |
| ISH9 | MITE | **MITEHgi2** | 1 | - | 0 |
| ISH9 | MITE | <u>MITEHgi5</u> | 1 | - | 0 |
| ISH14 | ISH | - | 0 | ISHbo1 | 2 |
| ISH14 | ISH | **ISHgi16** | 1 | - | 0 |
| ISH14 | ISH | HfIRS8 | 1 | - | 0 |
| ISH14 | ISH | HgiIRS46 | 1 | - | 0 |
| ISH7 | ISH | <u>ISHgi17</u> | 1 | - | 0 |
| ISHwa16 | ISH | - | 0 | HgiIRS21 | 1 |
| ISHwa16 | ISH | HgiIRS48 | 1 | - | 0 |
| ISHwa16 | ISH | HgiIRS49 | 1 | - | 0 |
| IS607 | ISH | **ISHgi6** | 1 | - | 0 |
| IS605 | ISH | **ISHgi7** | 2 | - | 0 |
| IS605 | ISH | **ISHgi15** | 1 | - | 0 |
| IS1341 | ISH | - | 0 | **ISHgi2** | 1 |
| IS1341 | ISH | - | 0 | **ISHgi3** | 1 |
| IS1341 | ISH | ISHvo18 | 1 | - | 0 |
| IS1341 | ISH | **ISHgi8** | 1 | - | 0 |
| IS1341 | ISH | <u>ISHgi9</u> | 1 | - | 0 |

| | | | | | |
|---|---|---|---|---|---|
| IS1341 | ISH | **ISHgi10** | 1 | - | 0 |
| IS1341 | ISH | **ISHgi11** | 1 | - | 0 |
| IS1341 | ISH | **ISHgi12** | 1 | - | 0 |
| IS1341 | ISH | **ISHgi13** | 1 | - | 0 |
| IS1341 | ISH | **ISHgi14** | 1 | - | 0 |
| IS1341 | ISH | HgiIRS53 | 1 | - | 0 |
| IS1341 | MITE | <u>MITEHgi3</u> | 1 | - | 0 |
| IS1341 | MITE | <u>MITEHgi4</u> | 1 | - | 0 |
| IS1341 | solo | MITEHgi27 | 1 | MITEHgi27 | 1 |
| IS1341 | solo | - | 0 | MITEHgi28 | 1 |
| IS1341 | solo | - | 0 | MITEHgi29 | 1 |
| IS1341 | solo | - | 0 | MITEHgi30 | 1 |
| IS1341 | solo | MITEHgi31 | 1 | MITEHgi31 | 1 |
| IS1341 | solo | - | 0 | MITEHgi32 | 1 |
| IS1341 | solo | - | 0 | MITEHgi33 | 1 |
| IS1341 | solo | MITEHgi34 | 1 | MITEHgi34 | 1 |
| IS1341 | solo | MITEHgi35 | 1 | MITEHgi35 | 1 |
| IS1341 | solo | MITEHgi61 | 1 | - | 0 |
| IS1341 | solo | MITEHgi62 | 1 | - | 0 |
| IS1341 | solo | MITEHgi63 | 1 | - | 0 |
| IS1341 | solo | MITEHgi64 | 1 | - | 0 |
| NmIRS44 | MITE | HgiIRS36 | 4 | HgiIRS36 | 3 |

**Table S7. Toxin/antitoxin systems of *Hfx. gibbonsii* LR2-5 and of *Hfx. volcanii* DS2$^T$.** Two types of toxins were found, those related to the VapC endonuclease, and those of the death-on-curing family. Orthologs in *Hfx. volcanii* or in *Hfx. gibbonsii* ARA6 are indicated, with % protein sequence identity (**seqid**) values given, with lower values highlighted in yellow. The lack of an ortholog is indicated by dash. Short adjacent genes are potential antitoxins. Such probable toxin/antitoxin pairs are highlighted in yellow (alternating between light and dark yellow). There are 5 unrelated types of short conserved hypothetical proteins (CHY1 to CHY5). A singlet second copy of the CHY1-type protein exists in strain LR2-5 and is more closely related to HVO_A0243 (highlighted in grey) than the copy that is adjacent to the toxin.

| ***Hfx. gibbonsii* LR2-5** | | | ***Hfx. volcanii* DS2$^T$** | | ***Hfx. gibbonsii* ARA6** | |
|---|---|---|---|---|---|---|
| **locus tag** | **len** | **class** | **locus tag** | **seqid (%)** | **locus tag** | **seqid (%)** |
| HfgLR_12920 | 131 | T;VapC(nf) | - | | - | |
| HfgLR_20875 | 132 | T;death_on_cur | HVO_A0244 | 87 | - | |
| HfgLR_20880 | 56 | A;CHY1 | HVO_A0243 | 85 | - | |
| HfgLR_20980 | 144 | T;VapC | - | | - | |
| HfgLR_20975 | 83 | A;VapB | - | | - | |
| HfgLR_21655 | 56 | A;CHY1 | HVO_A0243 | 94 | - | |
| HfgLR_22015 | 139 | T;VapC | - | | - | |
| HfgLR_22860 | 162 | T;death_on_cur | - | 53 | - | |
| HfgLR_22865 | 46 | A;CHY2 | - | | - | |
| HfgLR_23165 | 142 | T;VapC | HVO_A0188 | 95 | ABY42_18200 | 100 |
| HfgLR_23160 | 91 | A; CHY3 | HVO_A0189 | 93 | ABY42_18195 | 96 |
| - | 129 | T;death_on_cur | HVO_1841A | | - | |
| - | 41 | A; CHY4 | HVO_1842 | | - | |
| - | 160 | T;death_on_cur | HVO_A0391 | | - | |
| - | 45 | A; CHY5 | HVO_A0393 | | - | |

**Table S8**. **Restriction/modification systems and related endonucleases and methyltransferases of *Hfx. gibbonsii* LR2-5 and ARA6 and of *Hfx. volcanii* DS2$^{T}$.** For proteins with defined function, a gene name has been assigned (green highlighting) (*zim*: CTAG methyltransferase; *mrr*: Mrr family endonuclease; *rmeRMS*: type I restriction enzyme restriction/methylation/specificity subunit). For other proteins, only a general information is available (mrr-like: a protein, part of which is distantly related to mrr; RE domain: restriction endonuclease domain protein; DNA meth: probable DNA methyltransferase). For *Hfx. gibbonsii* ARA6, no attempt was made to identify corresponding genes that lack orthologs in strains LR2-5 and DS2$^{T}$.

| ***Hfx. gibbonsii* LR2-5** | | ***Hfx. volcanii* DS2$^{T}$** | | | ***Hfx. gibbonsii* ARA6** | |
|---|---|---|---|---|---|---|
| **locus tag** | **gene/class** | **locus tag** | **seqid (%)** | **gene/class** | **locus tag** | **seqid (%)** |
| HfgLR_04020 | *zim* | HVO_0794 | 91 | *zim* | ABY42_03790 | 99 |
| - | - | HVO_0682 | - | *mrr* | - | - |
| HfgLR_04065 | *mrr*-like | HVO_0803 | 85 | *mrr*-like | ABY42_03835 | 97 |
| - | - | HVO_1734A | - | RE domain | - | - |
| - | - | HVO_2269 | - | *rmeR* | ABY42_11400 | -/98 |
| - | - | HVO_2270 | - | *rmeM* | ABY42_11405 | -/82 |
| - | - | HVO_2271 | - | *rmeS* | - | - |
| - | - | HVO_2275 | - | DNA meth | - | - |
| - | - | HVO_A0079 | - | RE domain | - | - |
| - | - | HVO_A0006/A0237 | - | DNA meth | - | - |
| HfgLR_01370 | *rmeR1* | - | - | - | - | - |
| HfgLR_01375 | *rmeS1* | - | - | - | - | - |
| HfgLR_01380 | *rmeM1* | - | - | - | - | - |
| HfgLR_21960 | DNA meth | - | - | - | - | - |
| HfgLR_25220 | *rmeM3* | - | - | - | - | - |
| HfgLR_25230 | *rmeR2* | - | - | - | - | - |
| HfgLR_25235 | *rmeS2* | - | - | - | - | - |
| HfgLR_25240 | *rmeM2* | - | - | - | - | - |

**Table S9**. **The pilin-associated genes of *Hfx. gibbonsii* LR2-5 and of *Hfx. volcanii* DS2$^T$ with their orthologs/homologs in *Hfx. gibbonsii* ARA6.** For strains LR2-5 and DS2$^T$, gene names and locus tags are included and the listing is complete. Orthologs are listed in the same line, including the level of protein sequence identity. When two values are given for the protein from ARA6, the 1$^{st}$ shows sequence identity to the LR2-5 protein, the 2$^{nd}$ to that of the DS2$^T$ protein. The absence of an ortholog is indicated by a dash. For genes within a *pilBC* operon, the presence/absence of a type III signal sequence, as determined by FlaFind, is indicated. In one case (-/+), only the DS2$^T$ protein was FlaFind positive, while the LR2-5 protein was FlaFind negative. The PilBC proteins are not expected to have a type III signal sequence and thus were **n**ot **a**nalyzed (na). PilBC operons are marked by alternating light grey or dark grey shading.

| ***Hfx. gibbonsii* LR2-5** | | ***Hfx. volcanii* DS2$^T$** | | | | ***Hfx. gibbonsii* ARA6** | |
|---|---|---|---|---|---|---|---|
| **gene** | **locus tag** | **gene** | **locus tag** | **SpIII** | **seqid (%)** | **locus tag** | **seqid (%)** |
| *pibD* | HfgLR_15485 | *pibD* | HVO_2993 | na | 87 | ABY42_14925 | 99 |
| *pilB3* | HfgLR_05265 | *pilB3* | HVO_01034 | na | 95 | ABY42_05005 | 100 |
| *pilC3* | HfgLR_05260 | *pilC3* | HVO_01033 | na | 93 | ABY42_05000 | 99 |
| *pilA1* | HfgLR_04935 | *pilA1* | HVO_0972 | + | 51 | ABY42_04680 | 56/81 |
| *pilA2* | HfgLR_13125 | *pilA2* | HVO_2062 | - | 63 | ABY42_18020 | 65/55 |
| *pilA3* | HfgLR_11220 | *pilA3* | HVO_2450 | + | 85 | ABY42_12245 | 100 |
| *pilA4* | HfgLR_11215 | *pilA4* | HVO_2451 | + | 85 | ABY42_12250 | 100 |
| *pilA5* | HfgLR_24480 | *pilA5* | HVO_A0632 | + | 68 | ABY42_18015 | 71/74 |
| *pilA6* | HfgLR_24485 | *pilA6* | HVO_A0633 | + | 98 | - | - |
| DUF1628_1 | HfgLR_01515 | - | HVO_0308 | - | 90 | ABY42_01385 | 90 |
| DUF1628_2 | - | - | HVO_0875 | - | - | - | - |
| DUF1628_3 | HfgLR_09910 | - | HVO_2704 | + | 92 | - | - |
| - | - | *pilB1* | HVO_0620 | na | - | ABY42_02930 | -/93 |
| - | - | *pilC1* | HVO_0619 | na | - | ABY42_02925 | -/93 |
| - | - | - | HVO_0618 | + | - | ABY42_02920 | -/80 |
| - | - | - | HVO_0617 | - | - | ABY42_02905 | -/79 |
| - | - | - | HVO_0616 | + | - | ABY42_02910 | -/72 |
| - | - | - | HVO_0615 | - | - | ABY42_02915 | -/80 |

| | | | | | | | |
|---|---|---|---|---|---|---|---|
| - | - | - | HVO_0614 | + | - | ABY42_02920 | -/85 |
| - | HfgLR_03775 | - | HVO_0746 | - | 69 | ABY42_03555 | 96 |
| - | HfgLR_03780 | - | HVO_0747 | - | 86 | ABY42_03560 | 100 |
| *pilB2* | HfgLR_03785 | *pilB2* | HVO_0748 | na | 91 | ABY42_03565 | 98 |
| *pilC2* | HfgLR_03790 | *pilC2* | HVO_0749 | na | 92 | ABY42_03570 | 99 |
| - | HfgLR_03795 | - | HVO_0750 | - | 87 | ABY42_03575 | 98 |
| - | HfgLR_03800 | - | HVO_0751 | + | 88 | ABY42_03580 | 99 |
| - | HfgLR_03805 | - | HVO_0752 | - | 88 | ABY42_03585 | 98 |
| - | HfgLR_03810 | - | HVO_0753 | - | 85 | ABY42_03590 | 98 |
| - | HfgLR_03815 | - | HVO_0754 | - | 96 | ABY42_03595 | 100 |
| - | HfgLR_03820 | - | HVO_0755 | - | 94 | ABY42_03600 | 99 |
| *pilB4* | HfgLR_05935 | *pilB4* | HVO_1160 | na | 85 | ABY42_05670 | 99 |
| *pilC4* | HfgLR_05930 | *pilC4* | HVO_1159 | na | 95 | ABY42_05665 | 99 |
| - | HfgLR_05925 | - | HVO_1158 | + | 91 | ABY42_05660 | 96 |
| - | HfgLR_05920 | - | HVO_1157 | + | 94 | ABY42_05655 | 99 |
| - | HfgLR_05915 | - | HVO_1156 | + | 83 | ABY42_05650 | 98 |
| - | HfgLR_05910 | - | HVO_1155 | -/+ | 89 | ABY42_05645 | 98 |
| - | HfgLR_05905 | - | HVO_1154 | + | 89 | ABY42_05640 | 100 |
| *pilB5* | HfgLR_11565 | *pilB5* | HVO_2385 | na | 88 | ABY42_11900 | 99 |
| *pilC5* | HfgLR_11560 | *pilC5* | HVO_2386 | na | 94 | ABY42_11905 | 99 |
| - | HfgLR_11555 | - | HVO_2387 | - | 90 | ABY42_11910 | 99 |
| - | HfgLR_11550 | - | HVO_2388 | + | 87 | ABY42_11915 | 99 |

**Table S10. N-glycosylation sites in each LR2-5 PilA.** Predicted by https://www.hiv.lanl.gov/cgi-bin/GLYCOSITE/glycosite_main.cgi (Zhang *et al.*, 2004). Potential N-glycosylation sites of *Hfx. volcanii* are as described in (Esquivel *et al.*, 2016). As described, only a subset of sites exceed the threshold value of 0.5 (given in parenthesis). Only a subset of the N-glycosylation site is positionally conserved between orthologs. A dash indicates that no N-glycosylation sites are predicted for one of the orthologs.

| LR2-5 CDS | Name | N-glycosylation sites predicted in LR2-5 | N-glycosylation sites (*Hfx. volcanii*) | number of positionally conserved sites |
|---|---|---|---|---|
| HfgLR_04935 | PilA1 | 3 | 3(2) | 1 |
| HfgLR_11215 | PilA4 | 2 | 4(3) | 1 |
| HfgLR_11220 | PilA3 | 3 | 3(2) | 3 |
| HfgLR_13125 | PilA2 | - | 3 | - |
| HfgLR_24480 | PilA5 | 2 | - | - |
| HfgLR_24485 | PilA6 | 1 | 1 | 1 |

**Table S11**. **The genes involved in N-glycosylation or within the N-glycosylation cluster of *Hfx. volcanii* DS2$^T$ with orthologs from *Hfx. gibbonsii* LR2-5 and ARA6.** For all N-glycosylation genes of *Hfx. gibbonsii* LR2-5 see Table S9. Several transposons are integrated into the *Hfx. volcanii* N-glycosylation clusters but are not listed. For orthologs, % protein sequence identity (**seqid**) is indicated, with lower values highlighted in yellow. The lack of an ortholog is indicated by dash. Genes (**gene**) which are directly involved in N-gylcosylation have a gene name starting with "*agl*" and are highlighted dark green. Genes are classified, with a **class** that may be relevant for N-glycosylation being highlighted in light green. Classes are: glycosyltransferase (GT), nucleotidyltransferases which load a sugar onto an NTP (load), sugar modification enzymes (sug), oligosaccharyltransferases which thus transfer a glycan to a target protein (t2p), flippases (flip). Classes which are unrelated to N-glycosylation are highlighted in light grey: CHY: conserved hypothetical protein; AlkP: alkaline phosphatase core domain protein; CPxCG: small CPxCG-related zinc finger protein. The **task** represents schematically in which way the gene contributes to N-glycosylation. Tasks with "A" are involved in biosynthesis of the canonical N-glycan, tasks with "B" in the biosynthesis of the low-salt glycan. A or B followed by integer are the glycosyltransferases (class GT) which attach the corresponding sugar (1$^{st}$, 2$^{nd}$, etc) to the phosphorylated dolichol carrier. The term m() indicates enzymes which are involved in modification of the corresponding sugar. The task l() indicates enzymes which load a sugar to an NTP, and flip() indicates a flippase. The term flip(B4) indicates that the low-salt tetrasaccharide is moved across the membrane. For the genes flagged B1/2, it is yet unresolved which is responsible for attaching the 1$^{st}$ and which for the 2$^{nd}$ sugar.

| task | gene | class | *Hfx. volcanii* DS2$^T$ locus tag | *Hfx. gibbonsii* LR2-5 locus tag | seqid (%) | *Hfx. gibbonsii* ARA6 locus tag | seqid (%) |
|---|---|---|---|---|---|---|---|
| A5 | *aglD* | GT | HVO_0798 | HfgLR_04040 | 97 | ABY42_03810 | 97 |
| A1 | *aglJ* | GT | HVO_1517 | HfgLR_07690 | 90 | ABY42_07380 | 90 |
| m(A4) | *aglP* | sug | HVO_1522 | - | | - | |
| | *aglQ* | - | HVO_1523 | - | | - | |
| A4 | *aglE* | GT | HVO_1523A | - | | - | |
| flip(A5) | *aglR* | flip | HVO_1524 | - | | | |
| h(A5) | *aglS* | GT | HVO_1526 | - | | - | |
| l(A3) | *aglF* | load | HVO_1527 | HfgLR_13095 | 82 | ABY42_10470 | 82 |
| A3 | *aglI* | GT | HVO_1528 | - | | - | |
| A2 | *aglG* | GT | HVO_1529 | - | | - | |
| t2p(A4) | *aglB* | t2p | HVO_1530 | HfgLR_07755 | 76 | ABY42_07445 | 76 |
| m(A2) | *aglM* | sug | HVO_1531 | HfgLR_13100 | 64 | ABY42_10475 | 63 |

| | | | | | | |
|---|---|---|---|---|---|---|
| m(B1) | *agl7* | sug | HVO_2046 | - | - | |
| | | AlkP | HVO_2047 | - | - | |
| B3b | *agl9* | GT | HVO_2048 | - | - | |
| B4 | *agl10* | GT | HVO_2049 | - | - | |
| | | CHY | HVO_2052 | - | - | |
| B1/2 | *agl5* | GT | HVO_2053 | - | - | |
| flip(B) | *agl15* | flip | HVO_2055 | - | - | |
| m(B4) | *agl13* | sug | HVO_2056 | - | - | |
| l(B4) | *agl11* | load | HVO_2057 | - | - | |
| | | CPxCG | HVO_2057A | - | - | |
| m(B4) | *agl14* | sug | HVO_2058 | - | - | |
| m(B4) | *agl12* | sug | HVO_2059 | - | - | |
| m(B3) | *agl8* | sug | HVO_2060 | - | - | |
| B1/2 | *agl6* | GT | HVO_2061 | - | ABY42_10430 | 89 |

**Table S12. The genes involved in N-glycosylation or within the N-glycosylation cluster of *Hfx. gibbonsii* LR2-5 with orthologs of strain ARA6 and *Hfx. volcanii* DS2$^{T}$.** For all N-glycosylation genes of *Hfx. volcanii* DS2$^{T}$ see Table S8. Four groups of the genes are described in the text (**group**). Orthologs in *Hfx. volcanii* or in *Hfx. gibbonsii* ARA6 are indicated, with % protein sequence identity (**seqid**) values given, with lower values highlighted in yellow. The lack of an ortholog is indicated by dash. The **gene** name, if assigned to the *Hfx. volcanii* ortholog, is indicated. Genes are classified, with a **class** that may be relevant for N-glycosylation being highlighted in light green. Classes are: glycosyltransferase (GT), nucleotidyltransferases which load a sugar onto an NTP (load), sugar modification enzymes (sug), oligosaccharyltransferases which thus transfer a glycan to a target protein (t2p), flippases (flip). Classes which are unrelated to N-glycosylation (highlighted in light grey): CHY: conserved hypothetical protein; AlkP: alkaline phosphatase core domain protein; LpxA: LpxA domain protein. The task rfbX is not highlighted. These are annotated as "rfbX family transport protein" but should be considered flippase candidates.

| ***Hfx. gibbonsii* LR2-5** | | | ***Hfx. volcanii* DS2$^{T}$** | | | ***Hfx. gibbonsii* ARA6** | |
|---|---|---|---|---|---|---|---|
| **group** | **locus tag** | **gene** | **locus tag** | **seqid (%)** | **class** | **locus tag** | **seqid (%)** |
| - | HfgLR_04040 | *aglD* | HVO_0798 | 97 | GT | ABY42_03810 | 99 |
| 1 | HfgLR_07690 | *aglJ* | HVO_1517 | 90 | GT | ABY42_07380 | 100 |
| 1 | HfgLR_07695 | - | - | | GT | ABY42_07385 | 99 |
| 1 | HfgLR_07700 | - | - | | GT | ABY42_07390 | 99 |
| 1 | HfgLR_07705 | - | - | | rfbX | ABY42_07395 | 100 |
| 1 | HfgLR_07710 | - | - | | AlkP | ABY42_07400 | 99 |
| 1 | HfgLR_07715 | - | - | | AlkP | ABY42_07405 | 100 |
| 1 | HfgLR_07720 | - | - | | AlkP | ABY42_07410 | 99 |
| 1 | HfgLR_07725 | - | - | | AlkP | ABY42_07415 | 99 |
| 1 | HfgLR_07730 | - | - | | GT | ABY42_07420 | 99 |
| 1 | HfgLR_07735 | - | - | | LpxA | ABY42_07425 | 99 |
| 1 | HfgLR_07740 | - | - | | AlkP | ABY42_07430 | 100 |
| 1 | HfgLR_07745 | - | - | | AlkP | ABY42_07435 | 100 |
| 1 | HfgLR_07750 | - | - | | GT | ABY42_07440 | 100 |
| 1 | HfgLR_07755 | *aglB* | HVO_1530 | 76 | t2p | ABY42_07445 | 99 |
| - | HfgLR_12055 | - | - | | GT | ABY42_11260 | 100 |
| 2 | HfgLR_13050 | - | - | | AlkP | - | |
| 2 | HfgLR_13055 | - | - | | rfbX | - | |
| 2 | HfgLR_13060 | - | - | | GT | - | |

| | | | | | | | |
|---|---|---|---|---|---|---|---|
| 2 | HfgLR_13065 | - | - | | CHY | - | |
| 2 | HfgLR_13070 | - | - | | GT | - | |
| - | HfgLR_13075 | - | - | | GT | - | |
| - | HfgLR_13080 | - | - | | sug | - | |
| - | HfgLR_13085 | - | - | | sug | - | |
| 3 | HfgLR_13090 | - | - | | GT | - | |
| 3 | HfgLR_13095 | *aglF* | HVO_1527 | 82 | load | ABY42_10470 | 97 |
| 3 | HfgLR_13100 | *aglM* | HVO_1531 | 64 | sug | ABY42_10475 | 96 |
| 4 | HfgLR_21875 | - | - | | rfbX | - | |
| 4 | HfgLR_21890 | - | - | | AlkP | - | |
| - | HfgLR_23455 | - | - | | GT | ABY42_18470 | 100 |

#### Supplemental Video descriptions

**Video S1: Time-laps imaging of motile cells of *Haloferax gibbonsii* LR2-5.**

Cells from early-log growth phase ($OD_{600}$ 0.1 ) were prepared for time-laps-imaging after ~ 20 hours of growth at 42°C in 18% (w/V) salt CA-medium. 2 ml liquid culture were imaged at the base of a microscopy dish. Motile cells were imaged by phase-contrast microscopy, at 20 frames per second. Cells show swimming and tumbling behavior, some cells remain immotile.
